# NEUROEPITHELIAL BODIES AND TERMINAL BRONCHIOLES ARE NICHES FOR DISTINCTIVE CLUB CELLS THAT CAN REPAIR AIRWAYS FOLLOWING ACUTE NOTCH INHIBITION

**DOI:** 10.1101/2023.11.08.566345

**Authors:** Sai Manoz Lingamallu, Aditya Deshpande, Neenu Joy, Kirthana Ganeshan, Daniel Lafkas, Arjun Guha

## Abstract

Airway club cells (CCs) have the dual role of a secretory cell and a progenitor cell. Using pharmacological, genetic, and cell-ablation approaches we probe the role of canonical Notch signalling in the regulation of the regenerative capacity of CCs. We report that in response to its perturbation, different subpopulations of CCs adopt distinct fates. Upon acute inhibition of Notch, the majority transdifferentiate into multiciliated cells. However, a “variant” subpopulation (v-CCs), juxtaposed with Neuroepithelial Bodies (5-10%) and neighbouring bronchioalveolar duct junctions (>80%), does not. Instead, v-CCs transition into partially differentiated/lineage ambiguous states but can revert to a CC fate upon restoration of Notch signalling and repopulate the airways with CCs and multiciliated cells. Analysis of a v-CC lineage marker (*Uroplakin3a*), coupled with sequential Notch inhibition, reveals that differential responses of v-CCs to Notch inhibition are regulated by their cellular microenvironment. We propose that perturbations to Notch signalling may be a common consequence of airway injury and that microenvironmental signals diversify CCs to create a robust pool that can repair airways upon acute Notch inhibition.

## INTRODUCTION

The maintenance and repair of adult organs is dependent on specialised tissue-resident adult stem cells or on facultative stem cells that become activated in some manner by injury. The balance of these reparative cell populations varies across tissue types and even across different regions within a tissue. Tissues that turn over rapidly such as the gut, the skin, and the blood, are predominantly dependent on specialised stem cells whereas tissues that turn over more slowly, such as the epithelia comprising the lung and liver, may utilize both, with facultative stem cells making a significant contribution^1^. In this study, we probe how a facultative stem cell type that maintains and repairs the airway epithelium, the secretory club cell (CC, previously called Clara cell), is regulated.

CCs together with multiciliated cells are the major cell types that comprise the airway epithelium. These cells constitute the mucociliary apparatus that keeps the airways clear of inhaled toxicants^2,3^. In the murine lung, all CCs express the secretoglobin Scgb1a1/CC10, the definitive marker for this cell type^4^. In the human lung, although secretory cells are distributed throughout the airways, secretory cells expressing Scgb1a1 are located in the smaller airways^5^. In addition to mucociliary clearance, CCs contribute to xenobiotic metabolism, regulate inflammatory responses and serve as facultative stem cells^4–10^. They are typically quiescent and become activated in response to airway injury to self-renew and generate post-mitotic multiciliated cells^11,12^. Furthermore, CCs may also be mobilised in response to certain types of alveolar injury to generate both alveolar type 1 and alveolar type 2 cells^12,13^.

Recent studies have indicated that Notch has an important role in the regulation of CCs in the adult lung. The canonical Notch pathway, an evolutionarily conserved mechanism of juxtacrine signalling, becomes activated when transmembrane Notch receptors expressed on signal-receiving cells (Notch1, 2,3 and 4 in mammals) are bound by transmembrane ligands on signal-sending cells (Jagged1 and 2, Delta-like 1,3 and 4). Receptor-ligand engagement leads to the γ-secretase-dependent cleavage of the intracellular domain of the Notch receptor (NICD). Once cleaved, NICD translocates to the nucleus, binds to a preassembled transcriptional complex comprising the transcription factor RBPJκ and activates expression of Notch target genes^14,15^. Pertinently, Lafkas and colleagues showed that antibody-based inhibition of Notch in the adult lung in mice, via administration of Anti Jagged1 and Anti Jagged2 antibodies, induced transdifferentiation of the vast majority of the CCs into multiciliated cells in a matter of days^16^. The study also reported that rare CCs, enriched at certain anatomically defined tissue microenvironments in the airways, did not transdifferentiate into multiciliated cells. Dispersed within the airway epithelium, mostly at airway branchpoints, are rare groups of neuroendocrine cells called neuroepithelial bodies (NEBs)^17–19^. The CCs that escaped transdifferentiation appeared to be enriched at NEBs and near junctions of airways and alveoli (Bronchioalveolar Duct Junctions (BADJs)). These CCs were instrumental in restoring the balance of cell types post Notch inhibition.

Although the role for Notch in the regulation of the CC fate in the adult is a relatively recent discovery, several studies have previously elucidated a “developmental” role for Notch signalling in the specification of airway cell fate. Conditional deletion of RBPJκ in the fetal airway epithelium in mice leads to the complete loss of secretory cells and to supernumerary multiciliated and neuroendocrine cells^20–22^. Notch2 has been identified as the predominant receptor that regulates the balance of cell fates in the developing lung^22^. This “developmental” role for Notch signalling is also recapitulated in the adult lung in response to injury^21,23^. Basal cells, specialised airway stem cells that are located in the proximal airways of the mouse lung and more broadly in the human lung^24,25^, require Notch signalling to generate both secretory and multiciliated cells^26^. Basal cells lacking Notch signalling proliferate but only generate multiciliated cells^27^. Taken together, the studies on Notch signalling in the fetal and adult lung show that the pathway is required both for the specification of CCs and, potentially, for the maintenance of CC fate post specification. However, the occurrence of CCs in the adult lung that escape transdifferentiation into multiciliated cells upon antibody-mediated inhibition of Notch suggests that Notch signalling may be dispensable for the maintenance of these CCs.

We are interested in the mechanisms that regulate facultative stem cells during lung maintenance and repair. Thus, in an effort to delineate the role of canonical Notch signalling in the regulation of CCs, we adopted a systematic, multi-pronged approach for perturbing Notch signalling in the lung. Our pharmacological, genetic and cell ablation-based approaches revealed that Notch signalling is required for the stabilisation of all CCs. Importantly, we found that the rare cells at NEBs and at terminal bronchioles (∼0-200μm from the BADJs), that do not transdifferentiate into multiciliated cells in response to Notch signal inhibiting antibody treatment, lose secretory character and transition into lineage ambiguous states from which they can revert to a CC fate. The findings presented here elucidate the role of the Notch signalling pathway in the regulation of CCs in the adult lung and bring to light the mechanisms that, in concert, comprise a robust facultative stem cell pool for airway maintenance and repair.

## RESULTS

### Impact of antibody mediated inhibition of Notch signalling on airway CCs

In the homeostatic airway epithelium, Notch signalling has been implicated in the maintenance of the CC fate. Inhibition of Notch, using antibody antagonists that bind the Notch ligands-Jagged1/2 and prevent activation, induced transdifferentiation of the vast majority of CCs into multiciliated cells^16^. Under these conditions, rare CCs, associated with NEBs and enriched at terminal bronchioles, retained CC10 expression and seemingly escaped transdifferentiation. Although the implication of this study was clear, several outstanding questions remained. First, is canonical Notch signalling necessary for the maintenance of CC fate? Second, with regards to the CCs that escaped, did these CCs lose Notch signalling but not transdifferentiate into multiciliated cells, or were they simply not exposed to sufficiently high doses of antibody? To address both these questions, we decided to probe Notch signalling in the regulation of the CC fate using a variety of approaches.

We first examined whether all CCs, including NEB-associated CCs, activate Notch signalling. Activation of the Notch receptor upon ligand binding leads to the γ-secretase-dependent cleavage of the intracellular domain and its nuclear translocation. Pertinently, Notch 2 is the principal receptor that regulates the CC fate in the adult lung, with some contribution from Notch 1^16^. To probe the status of Notch signalling in the airway we performed immunostaining for the Notch2 intracellular domain (N2ICD). We observed nuclear N2ICD in the nuclei of all CCs (Figure 1A, red arrowheads), including NEB-associated CCs (Fig 1B, red arrowheads). This indicates that Notch signalling is indeed active in all CCs. Next, we examined how inhibition of Notch signalling via Anti Notch2/Notch1 antibody treatment impacted Notch signalling and the CC fate. We injected mice (n=3) with either a non-specific IgG (Control, see methods) or a combination of Anti Notch2 and Anti Notch1 antibodies^16^, and harvested lungs at different time points for histological analysis (regimen shown in schematic Figure 1C). We examined N2ICD immunoreactivity in these lungs and observed a depletion of nuclear N2ICD in all CCs within 24-48 hours post Anti Notch2/Notch1 antibody treatment (Figure 1D-E). More specifically, administration of the antibodies led to the absence of nuclear N2ICD in the CCs within 24 hours post-injection (Figure 1D, red arrowheads). We inferred that all CCs activate Notch signalling and that treatment with Anti Notch2/Notch1 perturbs signalling.

**FIGURE 1.**
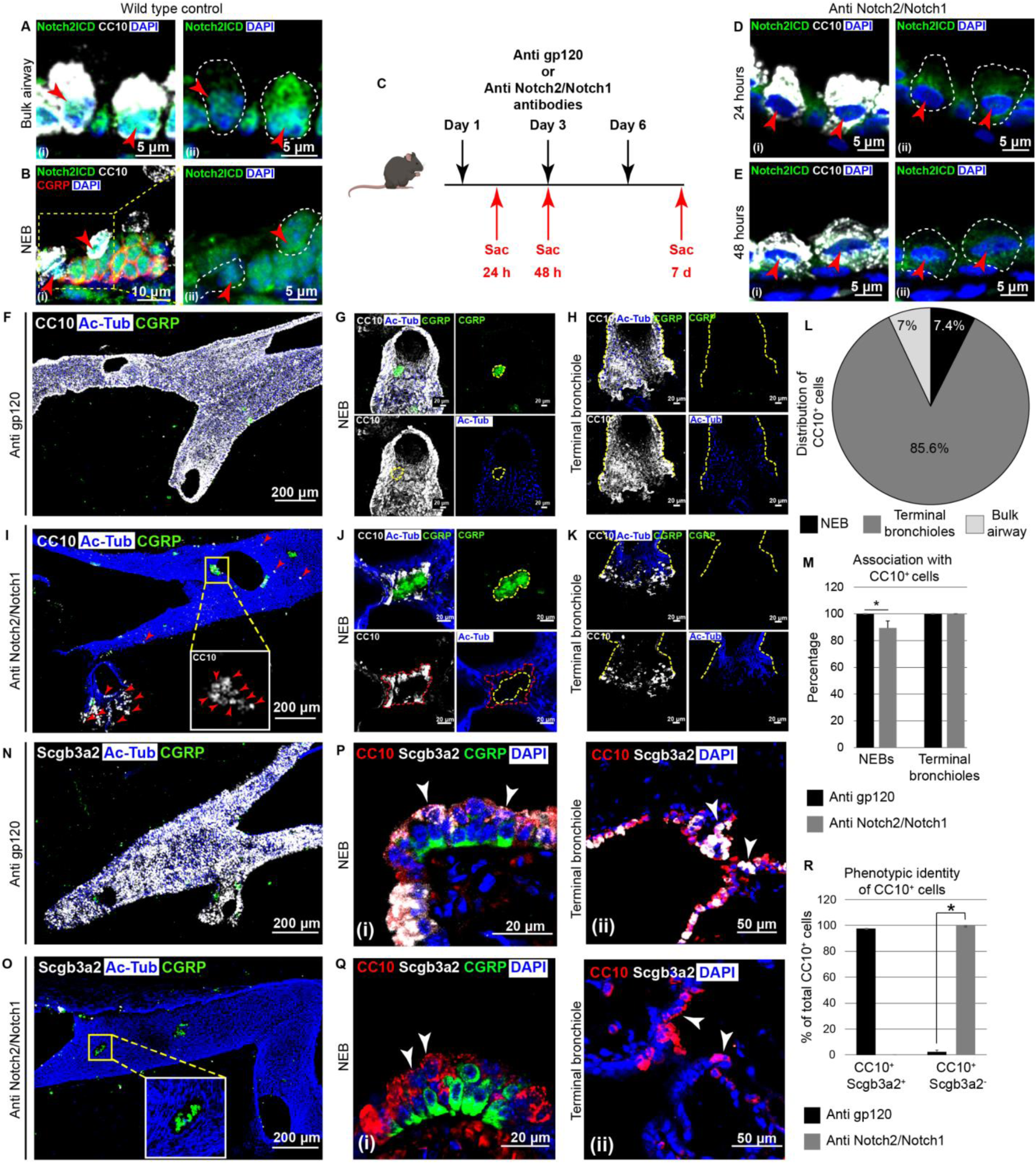
Antibody mediated inhibition of Notch2 and Notch1 impacts the fate of all airway CCs. (A-B) Notch signalling in CCs. Thin section of wild type lung showing the distribution of Notch2 intracellular domain (N2ICD, green) in bulk CCs (stained with Anti CC10, white, Ai, Aii) and in CCs surrounding Neuroepithelial bodies (NEBs, stained with Calcitonin gene-related peptide (CGRP) red, Bi, Bii). Red arrowheads point to nucleus. (C) Treatment regimen for inhibition of Notch2 and Notch1 using Control (Anti gp120) and Anti Notch2/Notch1 antibodies. (D-E) Notch signalling in CCs after antibody treatment. Localization of Notch2ICD (N2ICD, green) in bulk CCs (stained with Anti CC10, white), 24hr (Di, Dii) and 48hr (Ei, Eii) post antibody treatment. Red arrowheads point to nucleus. (F-K) Impact of antibody treatment on the CC fate. (F-H) Distribution of CCs (stained with Anti CC10/Scgb1a1, white), multiciliated cells (stained with Anti Acetylated tubulin (Ac-Tub), blue) and NEBs (stained with Anti CGRP, green) in thick sections (200μm) in control antibody (Anti gp120) treated lungs. (F) Low magnification view of the airways showing a salt-and-pepper pattern of CCs and multiciliated cells. (G-H) High magnification views of CCs around an NEB (demarcated with a yellow dashed line) (G) and at a terminal bronchiole (H). Note that NEBs are invariably surrounded by CCs and not multiciliated cells. (I-K) Distribution of CCs (stained with Anti CC10/Scgb1a1, white), multiciliated cells (stained with Anti Acetylated tubulin (Ac- Tub), blue) and NEBs (stained with Anti CGRP, green) in thick sections (200μm) from Anti Notch2/Notch1 antibody treated lungs. (I) Low magnification view of the airways showing CCs and multiciliated cells. Note depletion of CC10, red arrowheads indicate remaining CC10^+^ cells. Inset shows distribution of CC10^+^ cells around the boxed NEB. (J-K) High magnification views of CCs around a NEB (demarcated with a yellow dashed line) (J) and at a terminal bronchiole (K) after antibody treatment. The residual CC10^+^ cells do not express the multiciliated cell marker (red dashed line). (L) Distribution of CC10^+^ cells post Anti Notch2/Notch1 treatment. (M) Frequencies of NEBs and terminal bronchioles associated with CC10^+^ cells post antibody treatment. (N-Q) Impact of antibody treatment on Scgb3a2 expression in CCs/CC10^+^ cells. (N, P) Distribution of CCs (stained with Anti Scgb3a2, white), multiciliated cells (stained with Anti Acetylated tubulin (Ac-Tub), blue) and NEBs (stained with Anti CGRP, green) in thick sections (200μm) (N) and thin sections (5μm, DAPI - blue) (P) from control antibody treated lungs. Note the salt-and-pepper pattern of Scgb3a2-expressing cells and multiciliated cells in control (N). (P) White arrowheads in Pi and Pii indicate CCs double positive for CC10 and Scgb3a2 around an NEB (green) and at a terminal bronchiole respectively. Levels of Scgb3a2 can vary in the CC10^+^ cells in the terminal bronchioles. (O, Q) Distribution of CCs (stained with Anti Scgb3a2, white), multiciliated cells (stained with Anti Acetylated tubulin (Ac-Tub), blue) and NEBs (stained with Anti CGRP, green) in thick sections (200μm) (O) and thin sections (5μm, DAPI - blue) (Q) from Anti Notch2/Notch1 antibody treated lungs. Note the complete depletion of Scgb3a2 after antibody treatment (O,Q). White arrowheads in Qi and Qii indicate cells that are CC10^+^ but Scgb3a2^-^ around an NEB (green) and at a terminal bronchiole respectively. (R) Frequencies of CC10^+^Scgb3a2^+^ and CC10^+^Scgb3a2^-^ cells in antibody treated lungs. Data in pie chart are percentages from counts of the total number of cells from n=3 animals. Data in bar graphs represented as mean ± standard deviation, * denotes p <0.05, Student’s t-test. See also Figure S1.

To characterise the impact of Notch inhibition on CCs, we utilised two histological approaches coupled with immunostaining and confocal microscopy. First, to assess the geographic distribution of cells in the airways we performed immunostaining on 200μm thick precision-cut lung slices. Second, to assess the fates of individual cells we performed immunostaining on thin sections (5μm) from FFPE lungs. Both sets of assays were performed on at least 3-5 animals for each condition examined. By seven days post Notch inhibition, the airway epithelium was mostly devoid of CCs, as evidenced by the near absence of the CC marker CC10 and the pervasive distribution of multiciliated cell marker acetylated tubulin (Ac-Tub), (Figure 1I)). Surviving CC10^+^ cells were associated with NEBs (∼ 91% of NEBs were associated with CC10^+^ cells, Figure 1I, inset, 1J, 1L, 1M) and with terminal bronchioles ((∼0-200μm from the BADJs), Figure 1I, 1K, 1L, 1M). Aside from these locations, we also observed rare CC10^+^ cells scattered throughout the airway (Figure 1I, 1L). These results were consistent with the previous report that, upon Anti Notch2/Notch1 antibody treatment, bulk CCs transdifferentiate into multiciliated cells and that some CCs escaped this fate. Furthermore, we induced Scgb1a1^CreERTm/+^; Rosa^Tdtomato/+^ animals with tamoxifen to lineage tag all CCs, injected the animals with Anti Notch2/Notch1 antibodies and analysed the fates of Tdtomato expressing cells. While in the untreated lungs the Tdtomato^+^ cells were CC10^+^, in antibody-treated lungs the vast majority lacked CC10 and expressed markers of multiciliated cells (data not shown). In addition, we also noted that, under these conditions, Tdtomato^+^ cells surrounding NEBs and at terminal bronchioles, in both the control and post Anti Notch1/Notch2 antibody-treated lungs, remained CC10^+^ (data not shown).

In order to probe the impact of antibody-mediated inhibition of Notch on NEB-associated CCs, next we examined the expression of another CC marker - the secretoglobin Scgb3a2. CCs are typically double positive for CC10 and Scgb3a2 and express high levels of both proteins (Figure 1P). Whole mount sections from control lungs were stained with antisera to Scgb3a2 and Ac-Tub and imaged. Both markers were detected throughout the airways in a characteristic salt and pepper pattern (Figure 1F, 1N). Immunostaining of sections from Anti Notch2/Notch1 antibody-treated animals revealed a complete loss of Scgb3a2 expression throughout the airways; no Scgb3a2 was detected in cells around NEBs nor in any cells at terminal bronchioles (Figure 1O, 1Q, 1R). This suggested that treatment with the Notch antibody perturbs all CCs, including the CCs that resist transdifferentiation into a multiciliated fate.

To complement our study on Anti Notch2/Notch1 antibody-mediated Notch inhibition, we also treated the animals with Anti Jagged1/Jagged2 antibodies. Although both sets of antibodies inhibit Notch signalling with high efficacy, treatment with Anti Notch2/Notch1 antibodies is lethal and animals die within 7-8 days. In contrast, treatment with Anti Jagged1/Jagged2 antibodies is not lethal. Consequently, the Anti Jagged1/Jagged2 antibody treatment permits the analyses of cell fate long-term. Treatment with Anti Jagged1/Jagged2 antibodies induced a phenotype in the lung that was virtually indistinguishable from the Anti Notch2/Notch1 antibody treatment (Figure S1F-K). While bulk CCs transdifferentiated into multiciliated cells (Figure S1Gi), rare CCs, corralling NEBs (Figure S1Gii) and enriched at terminal bronchioles, appeared to resist transdifferentiation. These CCs retained CC10 but lost Scgb3a2 expression (Figure S1G, S1H, S1I). We examined lungs from Anti Jagged1/Jagged2-treated animals after an antibody washout period of 3-4 weeks. At this timepoint the airways were still largely devoid of CC10^+^ cells but, notably, the CC10^+^ cells around NEBs, and at terminal bronchioles, expressed both CC10^+^ and Scgb3a2^+^ (Figure S1L, S1M). We inferred that the loss of secretory character in CCs is a transient phenomenon and that they can revert to a CC fate upon restoration of Notch signalling.

In summary, the antibody-based inhibition of Notch led us to two unexpected findings. First, that the fate of all CCs is dependent on Notch signalling. Second, that a subpopulation of CCs, which we herewith refer to as variant-Club cells (v-CCs), lose secretory character upon Notch inhibition but resist transdifferentiation into a multiciliated fate.

### CCs that escape transdifferentiation post antibody-mediated inhibition of Notch signalling are lineage ambiguous post inhibition

Next we focussed our attention on v-CCs, both NEB and terminal bronchiole-associated populations, to Anti Notch2/Notch1 (and Anti-Jagged1/Jagged2) treatment. Immunostaining of thick sections from control lungs showed that NEBs are dense clusters of neuroepithelial cells and CCs (Figure 1G, 1N, Figure S1B). Although neuroepithelial cells and CCs are interspersed within NEBs, the peripheries of NEBs are invariably decorated with CCs; multiciliated cells are typically excluded from this microenvironment. Treatment with Notch-inhibitory antibodies induced the transdifferentiation of CCs in the bulk airways but not the CCs in the NEB microenvironment. A closer inspection of NEBs in these antibody-treated lungs revealed two interesting features. First, a few NEBs in the proximal airways (∼9% of the total number of NEBs, Figure 1M) completely lacked CC10^+^ cells and were surrounded by multiciliated cells (Figure S1Ei, S1J). Second, NEBs that remained associated with CC10^+^ cells (∼91%) had gaps in which we did not detect expression of any markers of CCs, neuroendocrine cells or multiciliated cells (Figure S1C - asterisks).

To examine the nature of these gaps, we stained thin sections with CC10, CGRP and FoxJ1. Immunostained sections from antibody-treated lungs revealed cells that did not express any of these markers (Figure S1Eii, S1K - asterisks). In order to identify these cells, next we stained sections with a suite of markers for other airway cell types – basal (K5), tuft (DCLK1), LNEP (Δp63), goblet (Muc5Ac) and ionocyte (Foxi1). These immunostainings revealed that the aforementioned cells did not express any of these markers either (Figure S2 C-G). We did find, however, that all cells were positive for Sox2 (Figure S2A) and TTF1 (Figure S2B) suggesting that these cells still retained lung, and airway, identity. Next we stained sections for markers of alveolar cells– alveolar type2 (SftpC/SPC, ABCA3) and alveolar type1 (Pdpn) (Figure 2 and Figure S2 H-I). Surprisingly, we observed SPC expression around many NEBs (Branchpoint NEBs – 80.34% ± 9.71%, Non-branchpoint NEBs – 65.18% ± 3.8%). In fact, a high frequency of the CCs juxtaposed to NEBs, (Figure 2C - red arrowheads, Figure 2D - white arrowheads, 2G) were SPC^+^. We also note that some cells in this microenvironment (∼4%) appeared to have lost the expression of CC10 (and Scgb3a2) and gained SPC expression (Figure 2D - asterisk, 2G). These CC10^-^SPC^+^ cells are likely to be cells that comprise the aforementioned gaps in the whole mount sections. Based on these data, we inferred that v- CCs juxtaposed to NEBs that do not transdifferentiate upon Anti Notch antibody treatment but lose secretory character (lack Scgb3a2 expression) become lineage ambiguous (lack CC10 expression and/or gain SPC expression).

**FIGURE 2.**
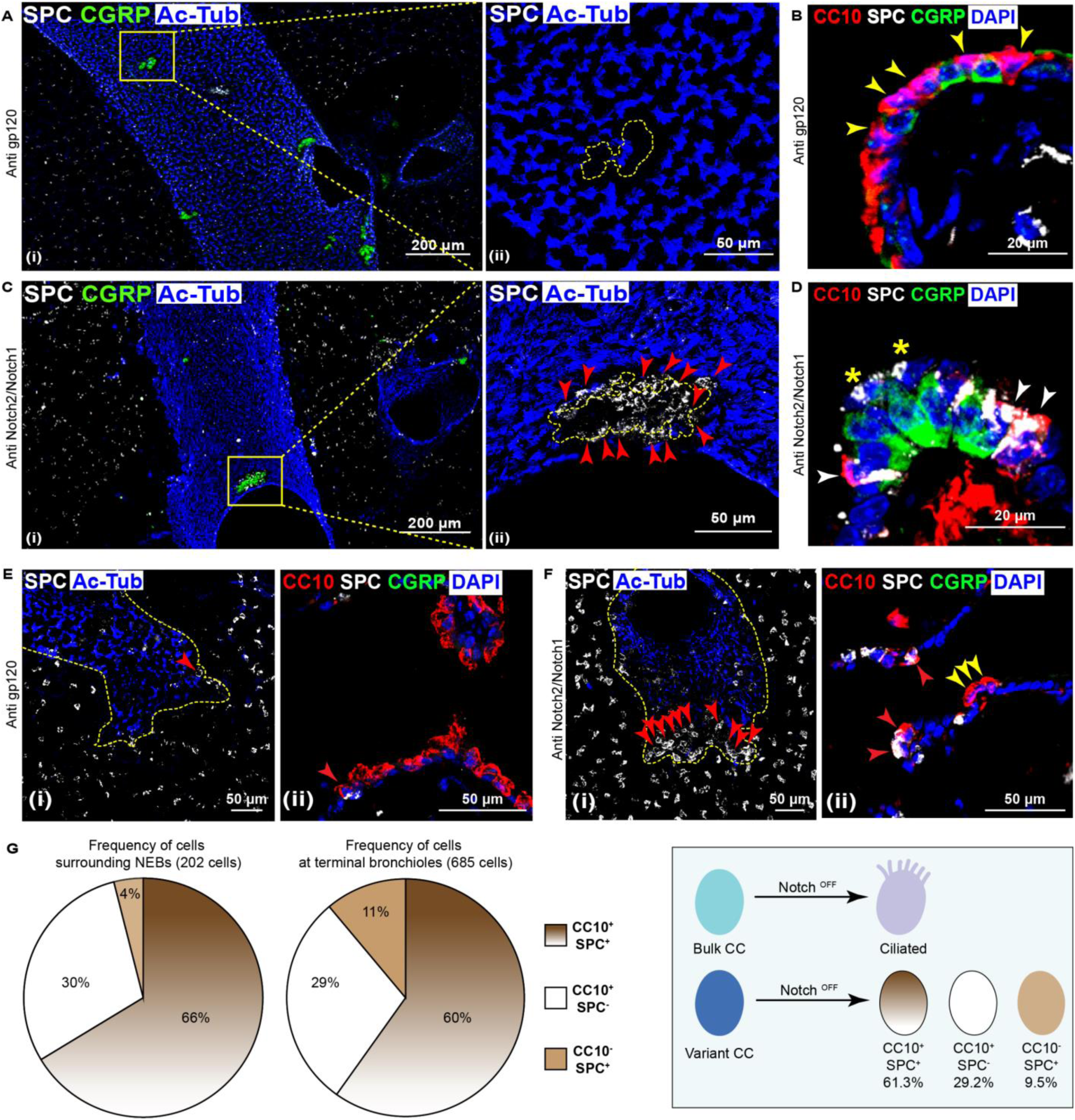
Many CCs surrounding NEBs and at terminal bronchioles become lineage ambiguous upon antibody-mediated inhibition of Notch2 and Notch1. (A-D) Phenotype of CCs around NEBs post antibody treatment. Distribution of alveolar type II marker (Surfactant protein C (SPC, white) around NEBs (green) in control antibody and Anti Notch2/Notch1 antibody treated lungs. (A) Low (Ai) and high (Aii) magnification views of a thick section from a control lung showing SPC expression around NEBs (demarcated by yellow dashed line). Note the lack of SPC expression around NEBs in control. (B) Thin section from a wild type lung stained for CC10/SPC/CGRP. Yellow arrowheads indicate CC10^+^SPC^-^ cells. (C) Low (Ci) and high (Cii) magnification views of a thick section from an Anti Notch2/Notch1 antibody treated lung showing SPC expression around NEBs (demarcated by yellow dashed line). Note SPC expression around NEBs (red arrowheads). (D) Thin section of Anti Notch2/Notch1 antibody treated lung stained for CC10/SPC/CGRP. White arrows indicate CC10^+^SPC^+^ cells. Yellow asterisks indicate CC10^-^SPC^+^ cells. (E-F) Phenotype of CCs at terminal bronchioles post antibody treatment. Distribution of alveolar type II marker (Surfactant protein C (SPC, white) at a terminal bronchiole in control (E) and Anti Notch2/Notch1 antibody treated lungs (F). (E) Distribution of alveolar type II marker (Surfactant protein C (SPC, white) at a terminal bronchiole in thick (Ei) and thin (Eii) sections of control lung. Red arrowheads in Ei, Eii indicate rare airway cells at BADJs that express SPC. (F) Distribution of alveolar type II marker (Surfactant protein C (SPC, white) at BADJs in thick (Fi) and thin (Fii) sections from Anti Notch2/Notch1 antibody treated lungs. Red arrowheads in Fi, Fii indicate many airway cells that express SPC. Note the presence of cells that co-express CC10 and SPC (Fii, red arrowheads) and only CC10 (Fii, yellow arrowheads). (G) Quantitative analysis of the phenotypes of cells around NEBs and at terminal bronchioles post Anti Notch2/Notch1 antibody treatment. Frequencies of cells that express CC10 or SPC in each environment is shown. Cartoon shows that cells expressing CC10 or SPC can be CC10^+^ SPC^+^, CC10^+^SPC^-^ or CC10^-^SPC^+^. Data in pie chart and cartoon are from a total of 887 cells across both microenvironments from n=3 animals. See also Figure S2, Figure S3.

v-CCs are enriched at both NEBs and terminal bronchioles. Terminal bronchioles are populated with equal frequencies of CCs and multiciliated cells. We noted that in response to Anti Notch2/Notch1 antibody treatment, a subset of the CCs at virtually every terminal bronchiole, all located 0-200 um from the BADJ, escaped transdifferentiation. Also, many of these CC10^+^ cells were SPC^+^ (Figure 2Fi, Fii - red arrowheads, 2G).

Taken together, the analysis of v-CCs post antibody-based Notch inhibition showed that they can adopt three distinct trajectories. First, they lose Scgb3a2 expression (Figure 1) and many among these turn on SPC expression. Second, some lose expression of secretory markers altogether and gain expression of SPC (Figure 2G). Third, a very small proportion of v-CCs may also differentiate into multiciliated cells (data not shown, also see next section). Treatment of lungs with Anti Jagged1/Jagged2 treatment recapitulated the phenotype observed post Anti Notch2/Notch1 treatment (Figure S3).

### Canonical Notch signalling in CCs is required to stabilise the CC fate

Having established that the CC fate is generally dependent on Notch receptors, next we probed whether the dependence reflects a role for canonical Notch signalling in regulating the CC fate. To test this, we inhibited Notch signalling using a γ-secretase inhibitor (Dibenzazepine, DBZ) (Figure S4). Treatment with DBZ recapitulated the phenotypes observed after Anti Notch2/Notch1 (and Anti Jagged1/Jagged2) antibody treatment, indicating a role for canonical Notch signalling in the maintenance of the CC fate. To directly test the role of canonical Notch signalling in CCs, we deleted the core transcription factor necessary for canonical Notch signalling, RBPJκ, in CCs using Scgb1a1^creERTm/+^; RBPJκ^flox/flox^ (Figure 3A) and examined the impact on CCs.

**FIGURE 3.**
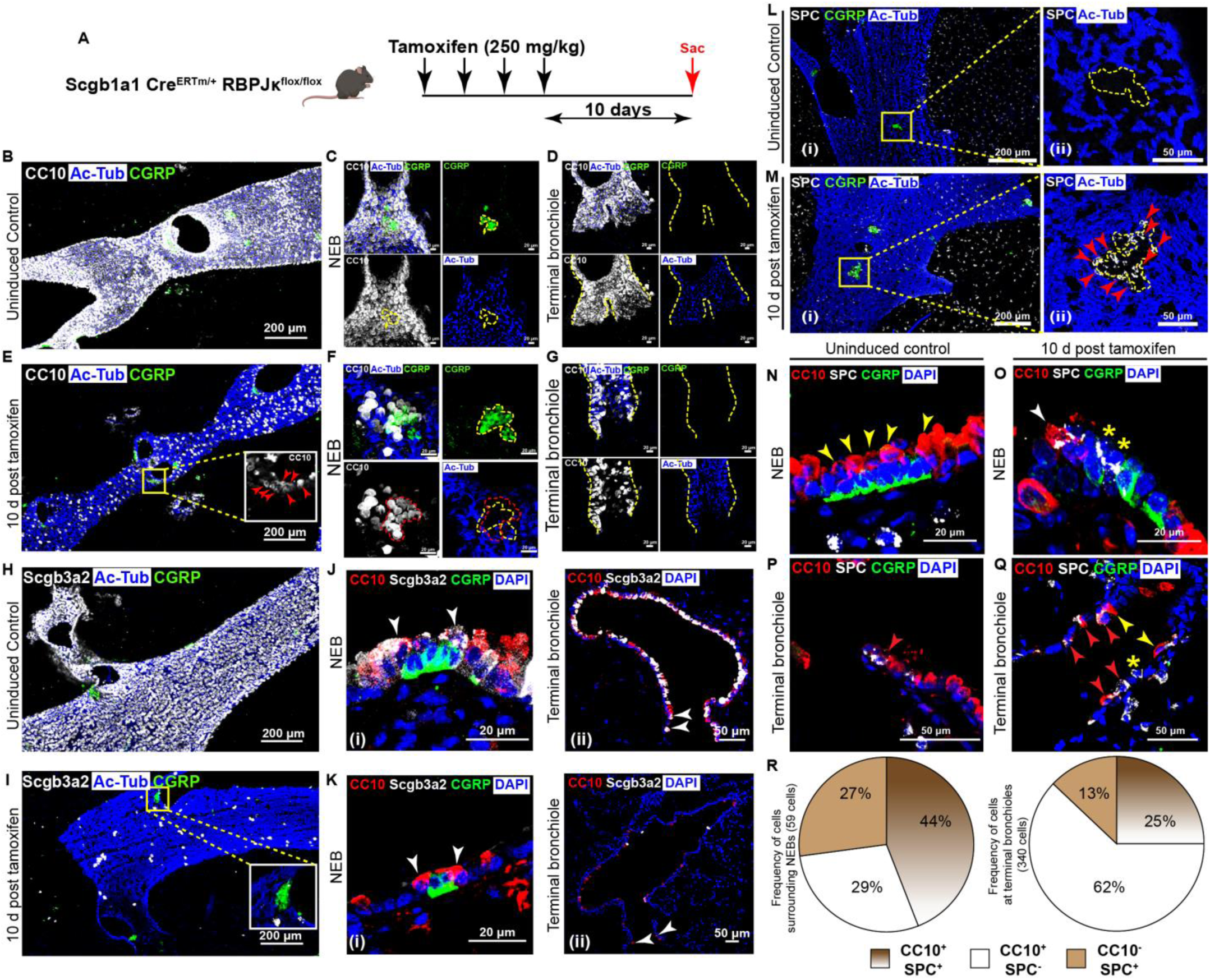
Genetic ablation of RBPJκ impacts the fate of all airway CCs. (A) Genetic strategy and tamoxifen regimen to conditionally deplete RBPJκ in CCs. (B-D) Distribution of CCs (stained with Anti CC10/Scgb1a1, white), multiciliated cells (stained with Anti Acetylated tubulin (Ac-Tub), blue) and NEBs (stained with Anti CGRP, green) in thick sections (200μm) from uninduced control lungs. (B) Low magnification view of the airways showing a salt-and- pepper pattern of CCs and multiciliated cells. (C-D) High magnification views of CCs around an NEB (demarcated with a yellow dashed line) (C) and at a terminal bronchiole (D). Note that NEBs are invariably surrounded by CCs and not multiciliated cells. (E-G) Distribution of CCs in RBPJκ-depleted airways (stained with Anti CC10/Scgb1a1, white), multiciliated cells (stained with Anti Acetylated tubulin (Ac-Tub), blue) and NEBs (stained with Anti CGRP, green) in thick sections (200μm) from tamoxifen treated lungs. (E) Low magnification view of the airways showing CCs and multiciliated cells. Note depletion of CC10. Inset shows distribution of CC10^+^ cells around the boxed NEB. (F-G) High magnification views of CCs around a NEB (demarcated with a yellow dashed line) (F) and at a terminal bronchiole (G) after RBPJκ-depletion. (H-K) Impact of RBPJκ-depletion on Scgb3a2 expression in CCs. (H, J) Distribution of CCs (stained with Anti Scgb3a2, white), multiciliated cells (stained with Anti Acetylated tubulin (Ac-Tub), blue) and NEBs (stained with Anti CGRP, green) in thick sections (200μm) (H) and thin sections (5μm, DAPI - blue) (J) from control lungs. Note the salt-and- pepper pattern of Scgb3a2-expressing cells and multiciliated cells in control (H). White arrowheads in (Ji) and (Jii) indicate CCs that co-express CC10 and Scgb3a2 around an NEB (green) and a terminal bronchiole respectively. Levels of Scgb3a2 vary in the CC10^+^ cells in the terminal bronchioles. (I, K) Distribution of CCs (stained with Anti Scgb3a2, white), multiciliated cells (stained with Anti Acetylated tubulin (Ac-Tub), blue) and NEBs (stained with Anti CGRP, green) in thick sections (200μm) (I) and thin sections (5μm, DAPI - blue) (K) in RBPJκ-depleted airways. White arrowheads in (Ki) and (Kii) indicate cells that are CC10^+^ but Scgb3a2^-^ around an NEB (green) and a terminal bronchiole respectively. (L-O) Phenotype of CCs around NEBs post RBPJκ-depletion. Distribution of alveolar type II marker (Surfactant protein C (SPC, white) around NEBs (green) in uninduced control and RBPJκ-depleted lungs. (L) Low (Li) and high (Lii) magnification views of a thick section from a control lung showing SPC expression around NEBs (demarcated by yellow dashed line). Note the lack of SPC expression around NEBs in control. (M) Low (Mi) and high (Mii) magnification views of a thick section from tamoxifen treated lung showing SPC expression around NEBs (demarcated by yellow dashed line). Note SPC expression around NEBs (red arrowheads). (N) Thin section from an uninduced control lung stained for CC10/SPC/CGRP. Yellow arrowheads indicate CC10^+^SPC^-^ cells. (O) Thin section from a RBPJκ-depleted lung stained for CC10/SPC/CGRP. White arrowhead indicates CC10^+^SPC^+^ cell. Yellow asterisks indicate CC10^-^SPC^+^ cells. (P, Q) Phenotype of CCs at a terminal bronchiole post RBPJκ-depletion. Distribution of alveolar type II marker (Surfactant protein C (SPC, white) at BADJs in thin sections of uninduced control (P) and RBPJκ-depleted (Q) airways. Red arrowheads indicate airway cells in the terminal bronchiole that express SPC. Note the presence of cells in Q that co-express CC10 and SPC (red arrowheads), only CC10 (yellow arrowheads) or only SPC (yellow asterisk). (R) Quantitative analysis of the phenotypes of cells around NEBs and at terminal bronchioles post RBPJκ-depletion. Frequencies of cells that express CC10 or SPC in each environment is shown. Data in pie chart are from a total of 399 cells across both microenvironments from n=3 animals. See also Figure S4, Figure S5, Figure S6.

To determine the efficiency of RBPJκ deletion in CCs in the airways of Scgb1a1^creERTm/+^; RBPJκ^flox/flox^ animals we immunostained thin sections from control and mutant animals (see methods) with anti RBPJκ antibody 10 d after tamoxifen induction (Figure S5C). RBPJκ expression is detected in all CCs in control lungs (Figure S5B). However, we did not detect any RBPJκ in most CCs in mutant lungs (∼94% of the CCs). This confirmed that RBPJκ had been deleted in most cells within 10 d post tamoxifen treatment (Figure S5C). Next we immunostained the same sections for CC markers to characterise the impact of RBPJκ ablation on the CC fate. There was a widespread reduction in the expression of both CC10 and Scgb3a2 and increased expression of acetylated tubulin in the airways of mutant lungs (Figure 3E, 3I). We inferred that the deletion of RBPJκ in CCs had resulted in the transdifferentiation of the majority of CCs into multiciliated cells.

A comparison of the frequencies of CC10^+^ cells in the RBPJκ mutant lungs with frequencies in lungs from animals treated with Anti Notch2/Notch1 (or Anti Jagged, Figure S1Gi) showed that CC10-expressing cells were more abundant in the former. Since most of the CC10^+^ cells in the RBPJk mutant lungs lacked expression of RBPJk, we inferred that these residual CC10^+^ cells were in transition to a multiciliated fate. To test this, we co-stained with FoxJ1, an early marker of commitment to a multiciliated fate. We found that many of the aforementioned CC10^+^ cells indeed co-expressed FoxJ1 and were likely in transition to a multiciliated fate (Figure S5Ci). Importantly, we observed that many NEBs and terminal bronchioles remained associated with CC10^+^ Scgb3a2^-^ cells (Figure 3F, 3G, 3K, S5D, S5E). These data establish that canonical Notch signalling in CCs is required for the stabilisation of the CC fate but that CCs respond differently to the loss of Notch signalling. To corroborate the cell-intrinsic requirement for canonical Notch signalling in CCs, we also deleted RBPJκ in multiciliated cells and found that deletion of RBPJκ in multiciliated cells did not change CC characteristics (Figure S6).

Next, we turned our attention to the cells that did not transdifferentiate into multiciliated cells around NEBs in tamoxifen induced Scgb1a1^creERTm/+^; RBPJκ^flox/flox^ lungs. The appearance of CC10^-^ Scgb3a2^-^ cells in this microenvironment led us to investigate the identity of these cells. We performed immunostainings for various airway markers as we had done post Anti Notch2/Notch1 treatment. We observed that many of the cells associated with NEBs expressed SPC after RBPJκ ablation (Figure 3M, 3O, quantified in 3R, ∼71%). This showed that post RBPJκ ablation there was an increase in both CC10^+^ SPC^+^ and CC10^-^SPC^+^ cells at NEBs (v-CCs did not begin expressing markers of other known cell types (data not shown)). Turning our attention to terminal bronchioles, we noted an increase in the frequencies of CC10^+^ SPC^+^ and CC10^-^SPC^+^ cells at these locations as well (Figure 3Q - red arrowheads, 3R). We infer that the vast majority of CCs transdifferentiate into multiciliated cells upon loss of canonical Notch signalling but a subpopulation resist transdifferentiation and transition into lineage ambiguous states instead.

Since, v-CCs reverted to a CC10^+^/Scgb3a2^+^ state post-acute Notch inhibition (1-month post Anti Jagged1/Jagged2 treatment (Figure S1L, M)), we wanted to analyse the phenotype of the v-CCs upon chronic Notch inhibition (1-month post RBPJκ ablation). At this time, we observed small clusters of CC10^+^/Scgb3a2^+^ CCs around NEBs, at terminal bronchioles and in the bulk airway. These CCs were RBPJκ^+^ indicating that these were cells that had escaped RBPJκ deletion (data not shown). Importantly, we detected far fewer CC10^+^ RBPJκ ^-^ cells in the bulk airways and at NEBs/terminal bronchioles. A closer inspection of the NEB microenvironment showed that the microenvironment now contained significantly higher frequencies of multiciliated cells (Figure S5J-L). We also quantified the frequencies of SPC^+^ cells at this timepoint to find that the frequencies of SPC^+^ cells were also significantly lower than at 10 d. Taken together, the data show that long-term or chronic inhibition of Notch signalling in v-CCs results in transdifferentiation into multiciliated cells or in cell death or both. This is in contrast to the effects of acute inhibition of Notch signalling, whereupon v-CCs revert to a CC fate.

### Ablation of multiciliated cells phenocopies aspects of Notch inhibition

The preceding experiments show that canonical Notch signalling in CCs is required for stabilising the CC fate and that CCs respond differently to the loss of Notch signalling. Multiciliated cells have been shown to express Notch ligands that activate Notch signalling in CCs^15^. This led us to hypothesise that loss of multiciliated cells would lead to a loss of Notch signalling in the CCs and would perturb the CC fate. Pertinently, the loss of multiciliated cells is known to occur in response to numerous environmental challenges^28–32^. To investigate this, we decided to ablate multiciliated cells in the adult lung by conditional expression of diphtheria toxin (DTA) in these cells. For this a FoxJ1^CreERT2^; Rosa^DTA/+^ strain was generated, induced with tamoxifen, and their lungs were analysed at various timepoints (Figure 4A).

**FIGURE 4.**
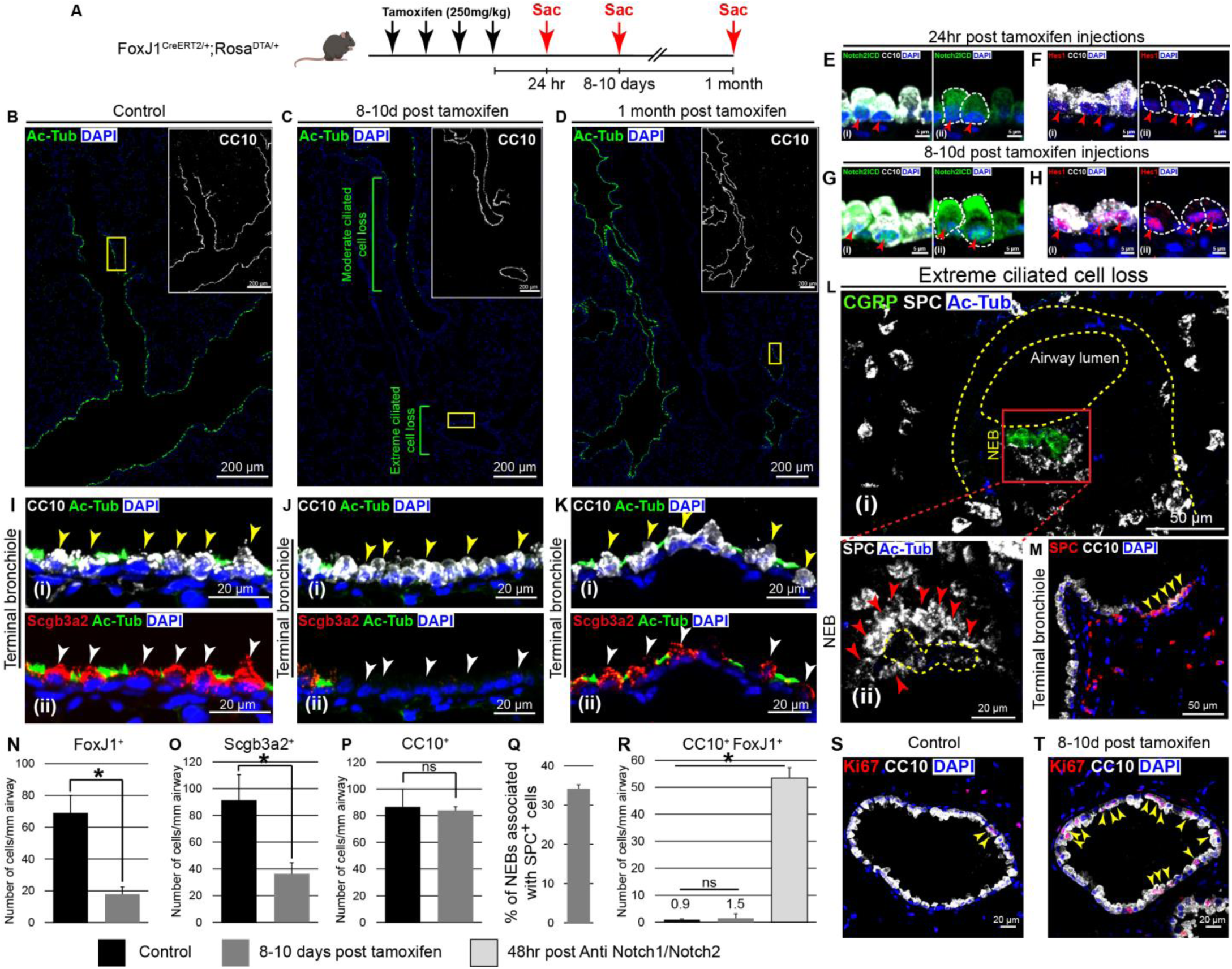
Ablation of multiciliated cells partially recapitulates the phenotypes of Notch pathway inhibition. (A-D) Impact of DTA induction in multiciliated cells on distributions of CCs and multiciliated cells. (A) Genetic strategy and tamoxifen regimen to induce DTA and conditionally ablate multiciliated cells. (B-D) Tiled images of the lung showing the distribution of the multiciliated cell marker - Acetylated tubulin (Ac-Tub, green) in control (B) and DTA-induced lungs at 8-10 d (C) and 1 month (D) post DTA induction. Airways were identified by counterstaining for CC10 (insets in B,C,D). High resolution images of the airways marked by yellow boxes in B,C,D are shown in I-K. (E-H) Impact of multiciliated cell ablation on Notch signalling in CCs. Thin section from tamoxifen treated FoxJ1^creERTm/+^; Rosa^DTA/+^ lungs showing the distribution of Notch2 intracellular domain (N2ICD, green) and Hes1 (red) in bulk CCs (stained with Anti CC10, white) in regions of multiciliated cell loss at 24hr (E,F) and 8-10 d post tamoxifen treatment (G,H). Red arrowheads indicate the nucleus. Note the absence of nuclear N2ICD and Hes1 at 24hr post tamoxifen treatment and their presence at 8-10 d post tamoxifen treatment. (I-K) Enlarged yellow boxed regions in B,C,D showing the impact of DTA induction on CCs in control (Ii, Iii) and DTA-expressing airways (J,K). Yellow arrowheads in Ii, Ji, Ki indicate the distribution of CC10. White arrowheads in Iii, Jii, Kii indicate the distribution of Scgb3a2 in the corresponding CCs. Note the depletion of Scb3a2 in CCs 8-10d post DTA induction and the restoration of Scgb3a2 by 1 month. (L,M) Phenotype of CCs surrounding NEBs and at terminal bronchioles post multiciliated cell ablation. (L) Distribution of alveolar type II marker (Surfactant protein C (SPC, white) around NEBs (green) in regions of lungs of extreme multiciliated cell ablation. Low magnification view of thick section (Li) and high magnification view of SPC expression around an NEB (demarcated by yellow dashed line, (Lii)) in the boxed area of Li. Note the gain of SPC expression around NEBs in regions of extreme multiciliated cell loss. (M) Thin section from lungs after multiciliated cell ablation stained for SPC/CC10. Yellow arrowheads indicate CC10^+^SPC^+^ cells at a terminal bronchiole. (N-P) Quantitative analysis of the frequencies of FoxJ1^+^ (N), Scgb3a2^+^ (O) and CC10^+^ (P) cells in the airways after multiciliated cell ablation. (Q) Quantitative analysis of the frequency of NEBs associated with SPC^+^ cells post multiciliated cell ablation. (R) Quantitative analysis of the frequencies of CC10^+^FoxJ1^+^ cells in the airways after multiciliated cell ablation. Note that there is no significant difference in the frequencies of CC10^+^FoxJ1^+^ cells in the control and multiciliated cell depleted airways indicating lack of transdifferentiation. Airways from 48hr post Anti Notch2/Anti Notch1 treatment were used to compare the frequencies of CC10^+^FoxJ1^+^ cells. (S,T) Phenotype of CCs post multiciliated cell ablation. Thin sections from wild type (S) and 8-10 d post tamoxifen treated (T) lungs stained for Ki67/CC10. Yellow arrowheads in (S) and (T) indicate Ki67^+^ CCs. Data represented as mean ± standard deviation, p <0.05 – Student’s t-test.

Tamoxifen induced DTA expression did indeed lead to the depletion of multiciliated cells in the airway 1-10 d after the last Tamoxifen injection (Figure 4C, 4N). Analysis of whole mount sections showed that the depletion of multiciliated cells was most acute in the distal airways, with some distal airways/terminal bronchioles having very few multiciliated cells. The reason for this mosaicism is currently unclear. Analysis of lungs 1 month post tamoxifen showed widespread distribution of multiciliated cells, indicating that the lungs had been repaired over this period (Figure 4D).

Next, we probed the impact of multiciliated cell ablation on Notch signalling in CCs by immunostaining for N2ICD and Hes1, a transcriptional target of Notch signalling, at 24 h and 8-10 d post tamoxifen. At 1 d, the levels of nuclear N2ICD and Hes1 were largely unperturbed in the larger airways (data not shown) but we noted a dramatic loss of both nuclear N2ICD (Figure 4E) and Hes1 (Figure 4F) in CCs in the distal airways. This decrease was most conspicuous in airway segments depleted of multiciliated cells. Interestingly, at 10 d post tamoxifen the levels of N2ICD and Hes1 in CCs were comparable to uninjured lungs throughout the airways, regardless of the abundance of multiciliated cells (Figure 4G, 4H). These data suggest that CCs have some mechanism by which they can compensate for the loss of Notch signalling induced by cell ablation. We then examined levels of Scgb3a2. The reduction in levels of nuclear N2ICD and Hes1 at 1 d were accompanied by the loss of Scgb3a2 expression; CC10 expression was unaffected (data not shown). At 10 d, levels of Scgb3a2 expression remained low (Figure 4Jii, 4O). At 1 month post tamoxifen, expression of N2ICD, Hes1, Scgb3a2 and CC10 were generally comparable to controls (Figure 4K, data not shown). We then probed the impact of multiciliated cell ablation on v-CCs. In regions of high multiciliated cell loss, v-CCs lost Scgb3a2 expression and gained the expression of SPC (Figure 4L - red arrowheads in 4Lii show SPC^+^ cells around an NEB, 4Q, 4M - yellow arrowheads show an expansion in SPC^+^ cells at a terminal bronchiole), a behaviour similar to conditions of Notch inhibition. Based on these findings we infer that multiciliated cell ablation can lead to a transient loss of Notch signalling in CCs and can partially phenocopy perturbations to Notch signalling itself.

CCs in the distal airways have been shown to proliferate and generate CCs and multiciliated cells during homeostasis and post-injury repair^4,12^. What was unclear from our above experiments is whether CCs directly transdifferentiate into multiciliated cells in response to multiciliated cell ablation. The characterization of CCs post Anti Notch/Jagged antibody treatment showed that the cells express CC10/Scgb3a2 initially, lose Scgb3a2 expression within 1-2 d post antibody treatment, and gain expression of FoxJ1, an early marker of commitment to the multiciliated fate, thereafter. At 2 d post antibody administration, there is a dramatic increase in the frequencies of cells co-expressing CC10 and FoxJ1 (Figure 4R). At 7 days post treatment, the frequencies of these double positive cells decrease and there is an increase in frequencies of CC10^-^ FoxJ1^+^ multiciliated cells. We hypothesised that if multiciliated cell ablation was inducing transdifferentiation of CCs, then there should be an increase in the frequencies of CC10-FoxJ1 double positive cells in airways depleted of multiciliated cells. To test this, we co-labelled sections from lungs isolated 1 d and 10 d post tamoxifen induction with antisera to CC10 and FoxJ1. There was no increase in the frequencies of double-positive cells, in airways depleted of multiciliated cells, at either of these timepoints (Figure 4R). Based on these findings we infer that there is little, if any, direct transdifferentiation of CCs into multiciliated cells post multiciliated cell ablation. Since there is efficient regeneration of multiciliated cells in 1 month following multiciliated ablation, we examined if CCs proliferate during this period. Sections immunostained with anti-ki67 antisera, a marker for cells that are mitotically active, showed that many CCs were mitotically active at 8-10 d post tamoxifen induction (Figure 4T). This suggests that the CCs proliferate and generate both CCs and multiciliated cells.

Taken together, these cell ablation experiments show that multiciliated cells contribute toward Notch signalling in CCs but that CCs respond differently to their ablation. We infer that CCs have compensatory mechanisms by which they are able to buffer the loss of Notch signalling induced by multiciliated cell ablation. It is plausible that this compensation prevents the direct transdifferentiation of CCs into multiciliated cells post ablation of the latter.

### Uroplakin 3a-expression distinguishes a subset of v-CCs and enables long-term tracing of daughter cells during repair post Notch inhibition

Analysis of lungs post Anti-Jagged1/Jagged2 treatment showed that the airways can restore the balance of cell types long-term^16^. Lineage analysis of Scgb1a1/CC10-expressing cells showed that CCs in distal airway contribute toward the repair^4,12^. In light of v-CC distribution detailed here, the question arises whether v-CCs at all locations contribute toward airway repair post antibody treatment. We have previously described a rare population of CCs in the adult lung, distinguished by Uroplakin3a (Upk3a) mRNA expression, that have a v-CC-like distribution^12^. More specifically, Upk3a-expressing cells are enriched around NEBs and terminal bronchioles. However, unlike v-CCs, nearly half the Upk3a-expressing cells, as illuminated by reporter gene expression, are juxtaposed with NEBs. Since only 5% of the v- CCs are located at NEBS, the distribution of Upk3a-expressing cells suggested that Upk3a expression tends to preferentially distinguish the NEB-associated v-CCs.

To probe the fate of v-CCs post Anti Jagged1/Jagged2 treatment long term, we utilised the previously described Upk3a^creERT/+^; Rosa^Tdtomato/+^ that selectively labels Upk3a-expressing CCs around NEBs and elsewhere (Figure 5B) (U-CCs). We treated tamoxifen induced Upk3a^creERT/+^; Rosa^Tdtomato/+^ animals with Anti Jagged1/Jagged2 antibodies and analysed the lungs at different time points (Figure 5A). 7 d post antibody treatment many Tdtomato^+^ cells around NEBs retain CC10 expression (Figure 5C – pink arrowheads, S7C, S7D – yellow arrowheads) and some cells among these express SPC (Figure S7E, S7F -yellow arrowheads). We infer that U-CCs are indeed a subset of v-CCs and estimate that they comprise 6.9 ± 2.1% (n=3 animals) of the total v-CC pool. Next, we analysed the fate of U- CCs in lungs three months and one year post Anti Jagged1/Jagged2 treatment. Analysis of whole mount sections 3 months post treatment revealed large patches of Tdtomato^+^ cells, comprised of both CCs and multiciliated cells (Figure 5D, 5E), radiating away from NEBS, and other locations in the airways including terminal bronchioles (Figure 5Eii). This suggests that U-CCs/v-CCs throughout the lung have the capacity to proliferate and restore the balance of cell types post Notch inhibition. Analysis of U-CC lineage 1 year post antibody treatment showed that Tdtomato^+^ CCs comprise 15 ± 5.1% of the total number of CCs formed.

**FIGURE 5.**
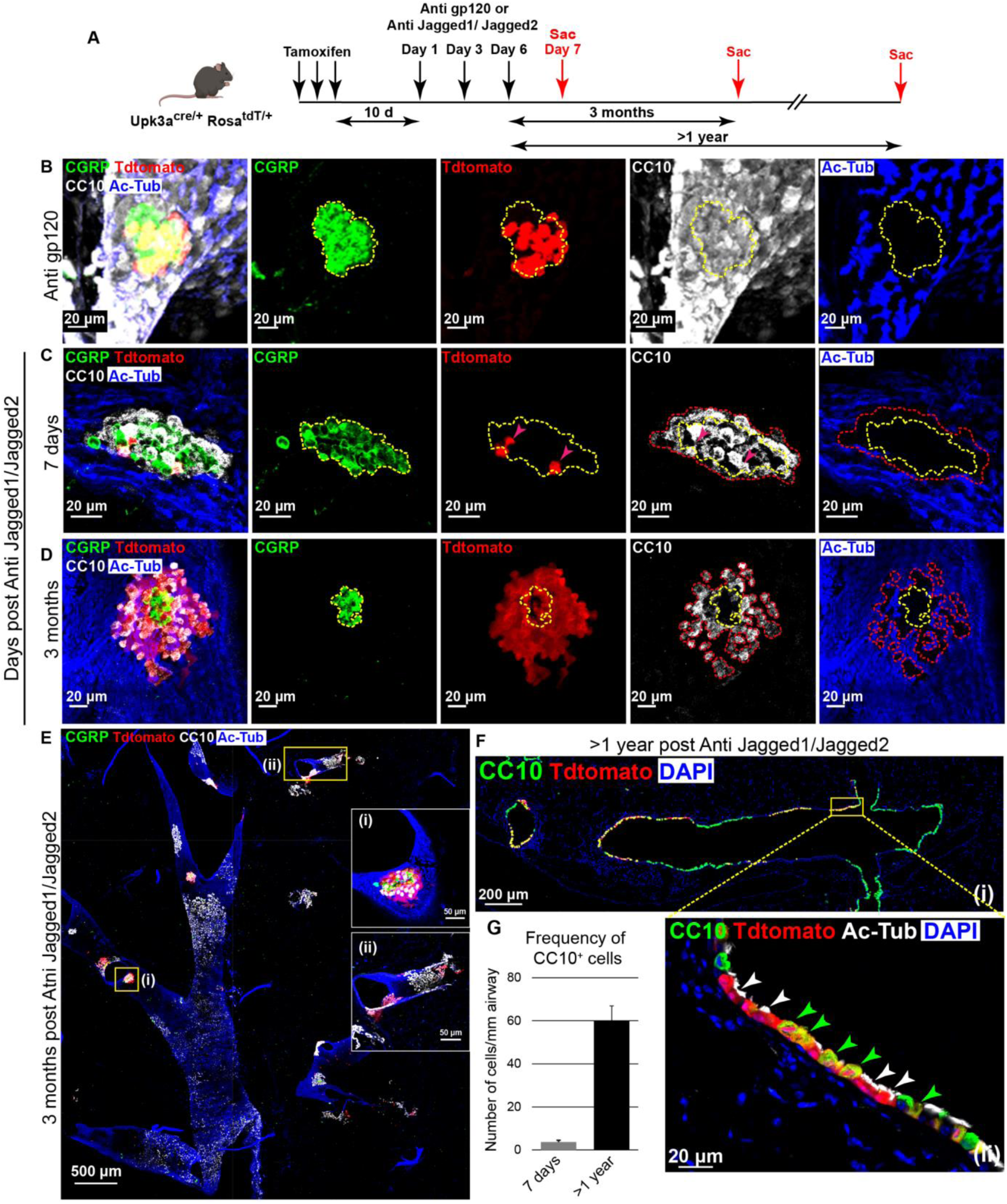
*Uroplakin3a^+^*CCs generate CCs and multiciliated cells post antibody mediated inhibition of Jagged1 and Jagged2 in the long-term. (A-F) Lineage analysis of U-CCs (v-CCs). (A) Treatment regimen for labelling U-CCs and inhibition of Notch signalling using antibodies in Upk3a^CreERTm/+^; Rosa^Tdtomato^ mice. (B-D) Distribution of CCs (stained with Anti CC10, white), U-CCs and their progeny (Tdtomato^+^), multiciliated cells (stained with Acetylated tubulin (Ac-Tub), blue) and NEBs (stained with Anti CGRP, green) in thick sections (200μm) of the lung. (B) Distribution of Tdtomato^+^ cells (U-CCs), CCs and multiciliated cells around an NEB (demarcated by yellow dashed line) in a control airway epithelium. (C) Distribution of CC10^+^ cells (demarcated by red dashed line) around a NEB (demarcated by yellow dashed line) 7 days post Anti Jagged1/Jagged2 treatment. Note that Tdtomato^+^ cells (U-CCs) (denoted by pink arrowheads) are CC10^+^. (D) Distribution of Tdtomato^+^ cells around an NEB (demarcated by yellow dashed line), 3 months post Anti Jagged1/Jagged2 treatment. Red dashed line demarcates CC10^+^ cells. (E) Low magnification view of a thick section (200μm) from 3 months post Anti Jagged1/Jagged2 treated lungs showing the distribution of Tdtomato derived patches in the airways. Insets are boxed regions indicating Tdtomato derived patches associated with an NEB (i) and at a terminal bronchiole (ii). (F) Thin sections from Anti Jagged1/Jagged2 treated lungs stained for CC10/Tdtomato post 1 year (Fi). (Fii) High resolution image of the airway marked by yellow box in Fi showing staining for CC10/Tdtomato/Ac-Tub. (G) Frequency of CC10^+^ cells post Anti Jagged1/Jagged2 treatment. Data represented as mean ± standard deviation. See also Figure S7.

### Cell-cell contact between v-CCs and neuroepithelial cells regulates the fate of v-CCs in the NEB microenvironment

The capacity of v-CCs to resist transdifferentiation in response to Notch inhibition and restore the balance of cell types post Notch inhibition, raises the question of how the fates of these cells are regulated. The spatial distribution of v-CCs, specifically the juxtaposition of the cells with neuroendocrine cells at NEBs, strongly suggests that cell-cell interactions are critical for the regulation of the v-CC fate. To probe this further, next we examined how the daughter cells derived from v-CCs at NEBs would respond to successive rounds of treatment with Anti Jagged1/Jagged2 antibodies. We hypothesized that if contact with neuroendocrine cells was necessary for the v-CC fate then only daughter cells in contact with NEBs would resist transdifferentiation.

We utilized the Upk3a^creERT/+^; Rosa^Tdtomato/+^ mice to probe the role of the microenvironment in regulation of v-CCs wherein we subjected these mice to two rounds of Anti Jagged1/Jagged2 treatment (Figure 6A). Upon a second round of Anti Jagged1/Jagged2 treatment, we observed that Tdtomato^+^ cells that were in direct contact with the NEBs retained CC10 expression (Figure 6B – white arrowheads) and did not transdifferentiate into multiciliated cells. However, the Tdtomato^+^ cells even one cell diameter away from the NEBs transdifferentiated into multiciliated cells (Figure 6B – yellow arrowheads).

**FIGURE 6.**
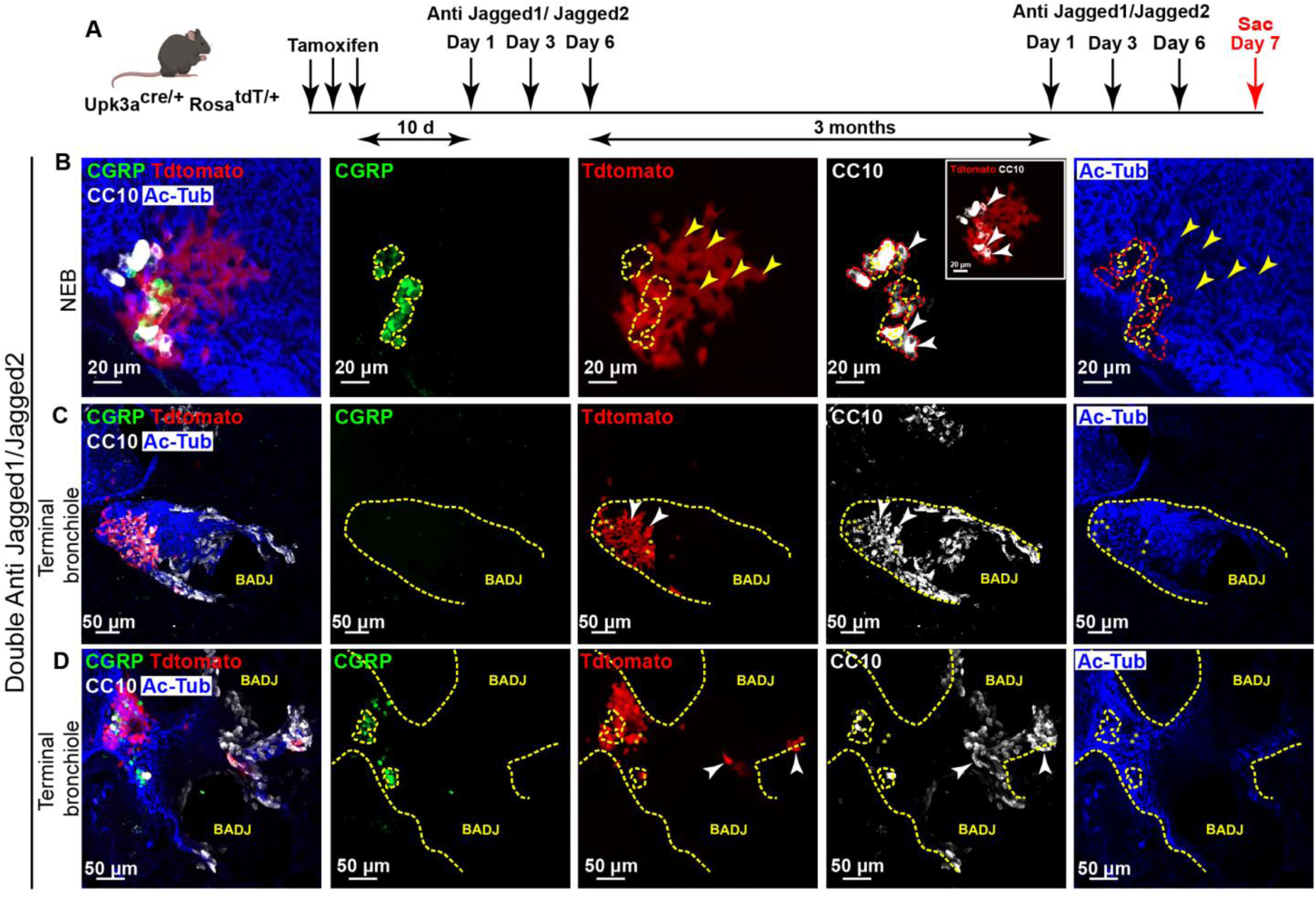
Responses of v-CCs to antibody-mediated inhibition of the Notch pathway are regulated by their contact with neuroepithelial cells in the NEB microenvironment. (A-D) Lineage analysis of U-CCs (v-CCs). (A) Treatment regimen for labelling U-CCs and inhibition of Notch signalling using antibodies in Upk3a^CreERTm/+^; Rosa^Tdtomato^ mice. (B) Distribution of Tdtomato^+^ cells around an NEB (demarcated by yellow dashed line), post double Anti Jagged1/Jagged2 treatment. Red dashed line demarcates CC10^+^ cells. Yellow arrowheads denote Tdtomato^+^ cells away from the NEB. Note that Tdtomato^+^ cells surrounding the NEB are CC10^+^ (white arrowheads). (C,D) Distribution of Tdtomato^+^ cells at terminal bronchioles (demarcated by yellow dashed line), post double Anti Jagged1/Jagged2 treatment. Yellow asterisks denote Tdtomato^+^ cells that are not CC10^+^. White arrowheads in (C,D) denote Tdtomato^+^CC10^+^ cells. Note that many Tdtomato^+^ cells close to the Bronchioalveolar Duct Junction (BADJ) are CC10^+^ in (C, white arrowheads) but those away from BADJ in (D, yellow asterisks) are not.

We also analysed the fates of rare patches of TdTomato^+^ cells at terminal bronchioles. All the patches of Tdtomato^+^ cells at terminal bronchioles 3 months post antibody treatment located within 200μm of the BADJs comprised both CCs and multiciliated cells. Pertinently, several Tdtomato^+^ cells in these patches expressing CC10 resisted transdifferentiation into multiciliated cells upon a second round of antibody treatment (Figure 6C, 6D, white arrowheads). Whether v-CCs in terminal bronchioles are also regulated by cell-cell contact remains to be established.

## DISCUSSION

The maintenance and repair of tissues is dependent both on “undifferentiated” stem cells and differentiated cells with the capacity for proliferation, self-renewal and multilineage differentiation (facultative stem cells)^1^. The epithelial lining of the respiratory tract utilizes both these types of cells. The aim of this study was to probe the regulation of facultative stem cells that maintain and repair the lower airways. Our study shows that all CCs are stabilized by Notch signalling, and that downregulation of Notch signalling leads to differential responses in CCs depending on their localization. We show that upon Notch inhibition, CCs either transition into a multiciliated fate or into less well defined, partial secretory/lineage-ambiguous states. Many of the cells in the latter pool are localized to NEBs but the vast majority are at BADJs (Figure 7A). These cells have the capacity to revert to a CC fate upon restoration of Notch signalling and to repopulate the airways with CCs and multiciliated cells. In the discussion that follows, we highlight aspects of regulation of facultative stem cells, in lung but also more generally, that have emerged from this work.

**Figure 7.**
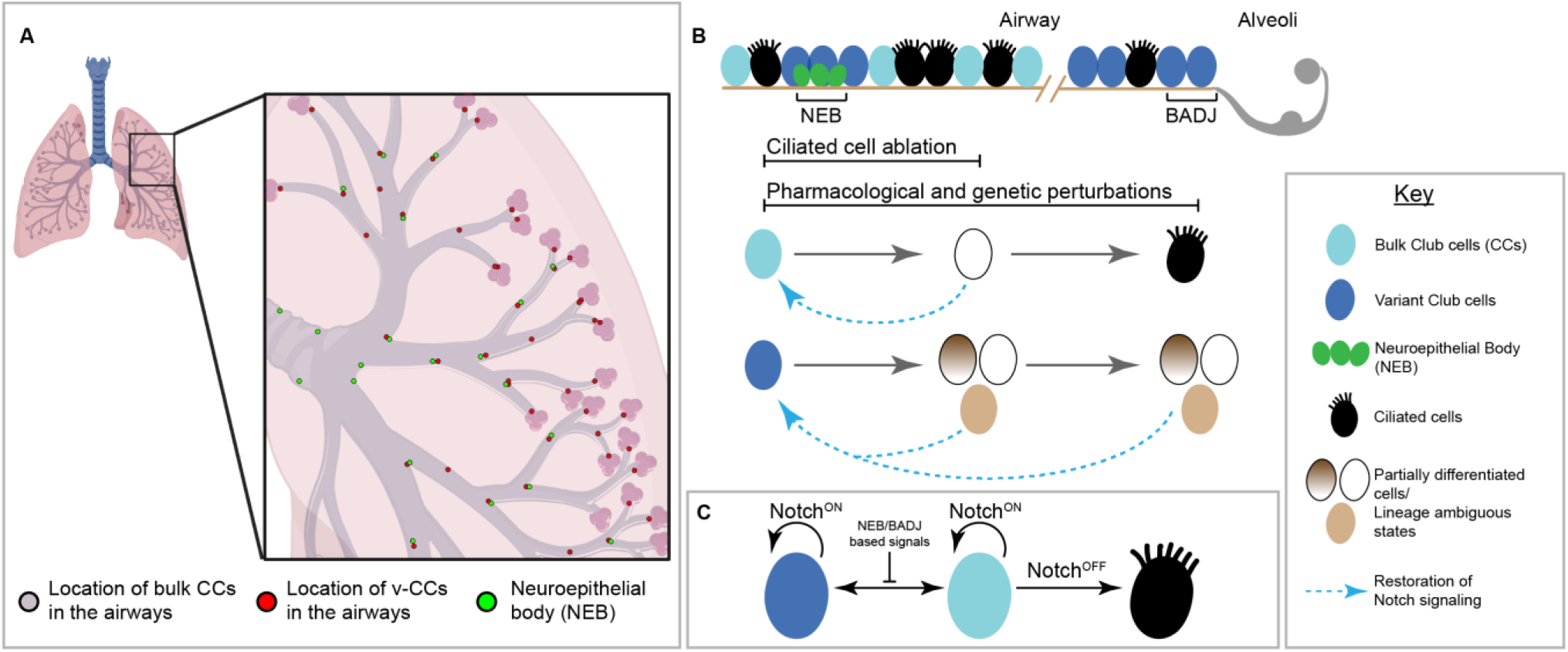
Microenvironmental signals regulate differences in cell state and lineage plasticity of club cells in the airways. (A) Graphical abstract showing the distribution of CCs, v-CCs and NEBs in the airways of the lung. (B) Responses of bulk CCs and v-CCs to perturbations to Notch signalling. (C) Regulatory framework of Notch and NEB/BADJ based signals in regulating the CC fate.

The mechanisms regulating undifferentiated tissue-resident stem cells have been described in considerable detail in the hematopoietic system, the musculature, and in barrier epithelial tissues like the skin and gut^33–41^. This body of work shows that the state of stem cells, and their responses, are orchestrated by their microenvironment or the “niche.” Stem cells are actively maintained in a dormant state by interactions with this microenvironment and are mobilized via changes in these interactions. Moreover, the analysis of stem and progenitor cell populations in the gut shows that there are several distinct cell populations that can contribute toward tissue renewal^42,43^. Our study on CCs in the lung suggests that the paradigms established for specialized stem cells are also likely to be relevant to facultative stem cells. The role for a juxtacrine signalling pathway like Notch in the stabilization of the CC fate, supports the concept that the fates of facultative stem cells are also regulated by the niches in which they reside (Figure 7C). Further, our findings implicate other short-range microenvironmental signals in the differential regulation of the fates of facultative stem cells across niches. The various approaches toward perturbing Notch signalling employed in this study show that the downregulation of Notch signalling in CCs is sufficient to induce cell fate transitions (Figure 7B). Although this suggests that the downregulation of Notch in CCs may be necessary for CCs to adopt alternate fates during CC-dependent regeneration, further studies are required to establish whether this indeed is the case.

The distinction between CCs and v-CCs described here raises the question about the adaptive value of this diversity. Two sets of observations shed light on this aspect. First, the multiciliated cell ablation experiment (Figure 4) suggests that the loss of multiciliated cells, particularly in the smaller airways, can lead to transient loss of Notch signalling in all CCs. This is consistent with the observation that multiciliated cells express high levels of Notch ligands Jag1 and Jag2^16,44^. We note that CCs can buffer this perturbation, compensate for the loss of multiciliated cells, and reinstate Notch signalling. We conclude that the ablation of multiciliated cells induces loss of secretory character in CCs but does not induce the transdifferentiation for this reason. This experiment does, however, bring to light the possibility that the airways, and the smaller airways in particular, are susceptible to the loss of Notch signalling. Similar perturbations in Notch signalling in the airways, albeit more modest, have also been reported in response to Bleomycin injury^44^. Thus, the diversification of CCs could be adaptive because they comprise a robust pool that can restore airways in response to any and all perturbations in Notch signalling in the lung. Second, we report that upon Notch inhibition, v-CCs transition into partial secretory/lineage ambiguous states. These cells may be CC10^+^Scgb3a2^-^SPC^-^ or CC10^+^Scgb3a2^-^SPC^+^ or CC10^-^Scgb3a2^-^SPC^+^ (Figure 2, 7B). Pertinently, CC10^+^SPC^+^ cells at BADJs have been reported by many groups, and these cells are thought to be dual-potential bronchioalveolar stem cells (BASCs)^45–47^ that can contribute to both airway and alveolar repair. An intriguing possibility that emerges from our study is that Notch inhibition (or change in some other signal) in v-CCs leads to their transition to BASCs. Here again, whether loss of Notch signalling is required for v-CC activation or additional signals can have this effect remains to be determined. The broader implication is that the distinction between bulk CCs and v-CCs could be adaptive because these cells have distinct capacities as facultative stem cells for lung repair. Consistent with this possibility is the finding that *Upk3a*-expressing CCs can contribute toward both airway and alveolar repair^12^.

The human lung, although similar in its overall architecture and make-up the murine lung, has some notable differences^2,25^. Are the findings presented here relevant to the human lung? Adult basal cell-derived 2D and 3D cultures can be induced to differentiate into secretory and multiciliated cells^24,48,49^. Treatment of these cultures with inhibitors of Notch signalling results in the loss of the secretory cells and to the generation of supernumerary multiciliated cells^50,51^ (KG, DL, unpublished observations). Cultures of adult nasal epithelial cells treated with a γ- secretase inhibitor exhibit a similar phenotype^52^. This suggests that the role of Notch as a stabilizer of the CC fate may be conserved in humans. The human lung has NEBs along the proximal-distal axis^53–55^. With regard to the diversity of CCs, we have shown previously that distal NEBs in the human lung may be associated with *Upk3a*-expressing CCs and comprise a niche for a distinctive subpopulation of CCs^12,56^. The transition zone from the airways to alveoli is markedly different in mice and humans. In contrast to the mouse lung, the human lung has a transition zone comprising respiratory bronchioles^57^. These are airway-like regions interspersed among alveolar regions. There is a large body of evidence, including the findings presented here, that BADJs in the murine lung harbour specialized CCs, including rare *Upk3a*- expressing cells and dual CC10-SPC-expressing BASCs, that can contribute toward alveolar repair^12,45–47^. Interestingly, two recent studies have pointed to distinctive secretory cell populations in the terminal and respiratory bronchioles of human lung^58,59^. One of these populations may be uniquely equipped to contribute toward alveolar repair^59^. Although there are significant differences in the architecture of murine and human airway:alveolar interface, the preponderance of v-CCs and BASCs at the BADJs in murine lungs, and the diversity of secretory cells in the terminal and respiratory bronchioles of the human lung, suggests that this interface constitutes a niche for a distinctive population of facultative stem cells across species.

## LIMITATIONS OF THE STUDY

While the role of Notch signalling in the regulation of secretory cells detailed here is likely to be conserved in humans, a role for human NEBs/BADJs as niches for distinctive secretory cells/facultative stem cells is not clear. A previous study from our group has established that *Upk3a* is detected around NEBs in the human lung, suggesting that the secretory cells in these microenvironments are distinct^12^. However, additional experiments are required to determine NEBs/BADJs are niches in the same capacity. Efforts to dissect the nature of the signals that regulate v-CCs, have been limited by the markers for these cells. The frequencies of *Upk3a*- expressing v-CCs are low and this limits the scope of bulk and single-cell RNA sequencing approaches to examine gene expression. Identification of additional markers for v-CCs, specifically the BADJ-associated population, should facilitate this analysis.

## ACKNOWLEDGEMENTS

We thank Dr. Mitsuru Morimoto (RIKEN, Japan) for providing us with the frozen embryos of the floxed RBP-J strain. We thank Dr. Rajesh Kumar Ladher (NCBS, India) for assisting us in procuring the frozen embryos of the floxed RBP-J strain. We thank the Central Imaging and Flow Cytometry Facility (CIFF) and the Animal Care and Resource Centre (ACRC) at the Bangalore Life Science Cluster (BLiSC) for their support. This work was supported by institutional core funds from inStem, DBT and a collaboration with Genentech (to A.G.). S.M.L. and A.D. are supported by a fellowship from the Department of Biotechnology (DBT), Government of India. N.J. is supported by a fellowship from Council of Scientific and Industrial Research (CSIR) (09/860(1273)/2019-EMR-I), Ministry of Science and Technology, Government of India. We thank Dr. Narmada Khare for critical reading of the manuscript.

## AUTHOR CONTRIBUTIONS

S.M.L., A.D., D.L. and A.G. designed the study and experiments. S.M.L., A.D. and N.J. performed the experiments and analysed the data. K.G. purified Anti Notch and Anti Jagged antibodies. S.M.L., A.D., D.L. and A.G. wrote the manuscript.

## STAR METHODS

All animal work reported here has been conducted in accordance with the necessary guidelines approved by the Institutional Animal Ethics Committee (IAEC) at inStem, Bengaluru. Detailed information regarding animal handling, mouse strains, Notch inhibition models, lineage analysis, histology, immunofluorescence, imaging, quantification of cell frequencies, flow cytometry and bulk-RNA sequencing, are provided in the Supplemental experimental procedures.

### Notch Inhibition Models

#### Antibody mediated Notch inhibition

B6/J, Upk3a^CreER/+^; Rosa26^Tdtomato/+^; or Scgb1a1^CreERTm/+^; Rosa26^Tdtomato/+^; animals (≥6 weeks of age) were injected intraperitoneally with Anti gp120 or Anti Notch1/Notch2 or Anti Jagged1/Jagged2 antibodies in PBS as indicated in the figures. Antibody concentrations used were as follows - Anti gp120 – 40mg/kg, Anti Notch 1 – 10mg/kg, Anti Notch 2 – 30mg/kg, Anti Jagged 1 – 20mg/kg and Anti Jagged 2 – 20mg/kg. Animals were sacrificed at the timepoints indicated in the figures.

#### Genetic ablation of RBPJκ in Club cells

Scgb1a1^CreERTm/+^; RBPJκ^flox/flox^ mice (≥6 weeks of age) were injected with Tamoxifen (Sigma, 250mg/kg body weight) four times on alternate days before sacrificing at the timepoints indicated in the figures.

#### Ablation of multiciliated cells

FoxJ1^CreERT2^; Rosa^DTA/+^ mice (≥6 weeks of age) were injected with Tamoxifen (Sigma, 250mg/kg body weight) four times on alternate days before sacrificing at the timepoints indicated in the figures.

### Genetic labelling of Club cells

Upk3a^CreER/+^; Rosa26^Tdtomato/+^; or Scgb1a1^CreERTm/+^; Rosa26^Tdtomato/+^ heterozygotes (≥6 weeks of age) were injected with Tmx (Sigma, 250mg/kg body weight) three times or once respectively on alternate days.

### Histology and Imaging

Analysis of cell populations was performed on vibratome (thick, (200 µm)) sections of paraformaldehyde-fixed, agarose-inflated lung tissues or thin (4-5 µm) sections of paraformaldehyde fixed, paraffin-embedded lung tissues. One lobe was fixed for two hours at room temperature and used to obtain thick sections while the other lobes were either fixed for 6-8 hours at room temperature or overnight at 4°C and processed for paraffin embedding. Stained sections were imaged either on Zeiss LSM-780 or Zeiss LSM-980 or Olympus FV3000 4/5 lasers, laser-scanning confocal microscopes.

### Quantification of cell frequencies

Cell frequencies were calculated from immunostained paraffin sections (4-5µm). Multi-channel images were acquired using confocal microscopes and cell frequencies were obtained by manual counting in images using Fiji (ImageJ). Student’s t-test was used for calculating the significance in the statistical analysis.

