## Supplemental Data for "NEUROEPITHELIAL BODIES AND TERMINAL BRONCHIOLES ARE NICHES FOR DISTINCTIVE CLUB CELLS THAT CAN REPAIR AIRWAYS FOLLOWING ACUTE NOTCH INHIBITION"

#### SUPPLEMENTARY FIGURE LEGENDS

**Figure S1. Impact of antibody mediated inhibition of Notch2/Notch1 and Jagged1/Jagged2 on CCs.** (A-E) Impact of Anti Notch2/Notch1 treatment on CCs. (A) Schematic showing the distribution of proximal NEBs, distal NEBs and v-CCs in the airways of the lung. (B,C) Distribution of CCs around an NEB. Distribution of CCs (stained with Anti CC10/Scgb1a1, white), multiciliated cells (stained with Anti Acetylated tubulin (Ac-Tub), blue) and NEBs (stained with Anti CGRP, green) in thick sections (200μm) in control (B) and Anti Notch2/Notch1 antibody treated (C) lungs. Red arrowheads denote $CC10^{+}$ cells. Note that $CC10^{+}$ cells surround the NEBs and some are interspersed with the neuroendocrine cells. Red asterisks in (C) denote regions devoid of CC10 and Ac-Tub around the NEB post Anti Notch2/Notch1 antibody treatment. Note that such regions are absent in the control. (D, E) Impact of antibody treatment on FoxJ1 expression in CCs/$CC10^{+}$ cells around NEBs. Thin sections from control antibody (D) and Anti Notch2/Notch1 antibody treated (E) lungs stained for CC10/FoxJ1/CGRP. White arrowheads in control antibody (D) and after Anti Notch2/Notch1 antibody treatment (E) denote $CC10^{+}$ cells. Yellow arrowheads in (Ei) denote $FoxJ1^{+}$ (multiciliated) cells after Anti Notch2/Notch1 antibody treatment. Note the occurrence of cells lacking CC, multiciliated and neuroendocrine markers after antibody treatment (denoted by asterisk in Eii) in distal NEBs. (F-M) Impact of Anti Jagged1/Jagged2 treatment on CCs. (F) Treatment regimen for inhibition of Jagged1 and Jagged2 using Anti Jagged1/Jagged2 antibodies. (G) Distribution of CCs (stained with Anti CC10/Scgb1a1, white), multiciliated cells (stained with Anti Acetylated tubulin (Ac-Tub), blue) and NEBs (stained with Anti CGRP, green) in thick sections (200μm) from Anti Jagged1/Jagged2 antibody treated lungs. (Gi) Low magnification view of the airways showing CCs and multiciliated cells. Note depletion of CC10, red arrowheads indicate remaining $CC10^{+}$ cells. (Gii) shows a high magnification view of the distribution of $CC10^{+}$ cells around the boxed NEB. (H, I) Impact of antibody treatment on Scgb3a2 expression in CCs/$CC10^{+}$ cells. Thin sections (5μm) from antibody treated lungs stained for CC10/Scgb3a2/CGRP (H) and CC10/Scgb3a2 (I). White arrowheads indicate cells that are $CC10^{+}$ but $Scgb3a2^{-}$ around NEBs (H) and at terminal bronchioles (I). (J, K) Impact of antibody treatment on FoxJ1 expression in CCs/$CC10^{+}$ cells around NEBs. Thin sections from Anti Jagged1/Jagged2 antibody treated lungs stained for CC10/FoxJ1/CGRP. Yellow arrowheads in (J) denote $FoxJ1^{+}$ (multiciliated) cells after Anti Jagged1/Jagged2 antibody treatment. White arrowheads in (K) denote $CC10^{+}$ cells. Note the occurrence of cells lacking CC, multiciliated and neuroendocrine markers after antibody treatment (denoted by asterisk in K) in distal NEBs. (L,M) Impact of Anti Jagged1/Jagged2 treatment on CC fate in the long term. Distribution of CCs (stained with Anti CC10 - red and Anti Scgb3a2 - white) around an NEB (stained with Anti CGRP, green, L) and at a terminal bronchiole (M) in thin sections (5μm) from 3-4 weeks post Anti Jagged1/Jagged2 antibody treated lungs. Note that CCs are double positive for CC10 and Scgb3a2 (yellow arrowheads).

**Figure S2. CCs surrounding NEBs retain airway identity but do not differentiate into other known airway or alveolar cell states upon antibody mediated inhibition of Notch2 and Notch1.** (A-I) Phenotypic identity of NEB associated CCs. (A-I) Thin sections from Anti Notch2/Notch1 treated lungs stained for CGRP (green), Ac-Tub or FoxJ1 (red) and markers of various cell types (white) of the airway epithelium - Sox2 (A), TTF1(B), basal cells (K5, C), tuft cells (DCLK1, D), Lineage negative epithelial progenitors (LNEPs, Δp63, E), goblet cells (Muc5Ac, F), ionocytes (Foxi1, G), alveolar type II cells (ABCA3, H) and alveolar type I cells (Pdpn, I). White arrowheads in the panels denote non-multiciliated, NEB associated cells. Insets represent positive control for immunostaining of the corresponding markers. Scale bar – 10 μm.

**Figure S3. CCs surrounding NEBs and at terminal bronchioles express an alveolar type II marker Surfactant protein C (SPC) upon antibody mediated inhibition of Jagged1 and Jagged2.** (A-B) Impact of Anti Jagged1/Jagged2 antibody treatment on CCs. Thin sections from Anti Jagged1/Jagged2 antibody treated lungs showing staining for CGRP/CC10/SPC around NEB (A) and at terminal bronchioles (B). White arrows indicate $CC10^{+}$ cells that gain SPC expression.

**Figure S4. Impact of small-molecule mediated (γ-secretase inhibitor - Dibenzazepine (DBZ)) inhibition of Notch signalling on CCs.** (A-H) Impact of DBZ treatment on CCs. (A) Treatment regimen of vehicle or DBZ. (B, C) Thin sections from vehicle control (B) and DBZ (C) treated lungs stained for FoxJ1/CC10. Note the depletion of CC10 expression after DBZ treatment (C). (D, E) Thin sections from vehicle control (D) and DBZ (E) treated lungs stained for CGRP/CC10. Note retention of CC10 expression in cells around NEBs after DBZ treatment (Ei). (Dii, Diii and Eii, Eiii) High magnification views of the boxed regions in Di and Ei respectively. Yellow dotted lines denote neuroendocrine cells. Yellow arrowheads denote $CC10^{+}$ cells. (F) Thin sections from DBZ treated lungs stained for CGRP/CC10/Scgb3a2. Yellow arrowheads denote $CC10^{+}$ cells around NEBs. Note the lack of Scgb3a2 expression after DBZ treatment. (G) Frequencies of $CC10^{+}Scgb3a2^{+}$ and $CC10^{+}Scgb3a2^{-}$ cells in vehicle and DBZ treated lungs. (H) Thin sections from DBZ treated lungs stained for

CGRP/CC10/SPC. Yellow arrowheads denote $CC10^+$ cells around an NEB. Note the gain of SPC expression in $CC10^+$ cells after DBZ treatment. Data represented as mean ± standard deviation, * denotes p <0.01, Student's t-test.

**Figure S5. Impact of RBPJκ depletion on CCs.** (A-C) Impact of tamoxifen treatment on RBPJκ expression. (A) Genetic strategy and tamoxifen treatment regimen in $Scgb1a1^{creERTm/+}$; $RBPJκ^{flox/flox}$ to conditionally ablate RBPJκ in CCs. Thin sections from uninduced control (B) and tamoxifen treated lungs (C) stained with RBPJκ /CC10/FoxJ1 (Bi, Ci) and RBPJκ /CC10 (Bii, Cii) . Note the loss of RBPJκ expression in CCs upon tamoxifen treatment. Yellow arrowheads denote the nucleus. (D-I) Impact of RBPJκ ablation on NEB associated CCs. (D) Frequencies of NEBs and terminal bronchioles associated with $CC10^+$ cells in lungs from uninduced control (black bars) and 10 days post RBPJκ ablation (grey bars). (E) Frequencies of $CC10^+Scgb3a2^+$ and $CC10^+Scgb3a2^-$ cells in lungs from uninduced control (black bars) and 10 days post RBPJκ ablation. (F) Distribution of CCs (stained with Anti CC10/Scgb1a1, white) and multiciliated cells (stained with Anti Acetylated tubulin (Ac-Tub), blue) in thick sections (200μm) around an NEB ((stained with Anti CGRP, green) from Figure 3F) from tamoxifen treated lungs. Red arrowheads denote $CC10^+$ cells. Note that $CC10^+$ cells surround the NEBs and some are interspersed with the neuroendocrine cells. Red asterisks in (F) denote regions devoid of CC10 and Ac-Tub around the NEB post tamoxifen treatment. (G-I) Thin sections from control (G) and tamoxifen treated (H, I) lungs stained for CC10/FoxJ1/CGRP. White arrowheads in control (G) and after tamoxifen treatment (I) denote $CC10^+$ cells. Yellow arrowheads in (H) denote $FoxJ1^+$ (multiciliated) cells after tamoxifen treatment. Note the occurrence of cells lacking CC, multiciliated and neuroendocrine markers after tamoxifen treatment (denoted by asterisk in I). (J-L) Impact of RBPJκ ablation on CCs in the long term. (J) Distribution of CCs (stained with Anti CC10/Scgb1a1, white), multiciliated cells (stained with Anti Acetylated tubulin (Ac-Tub), blue) and NEBs (stained with Anti CGRP, green) in thick sections (200μm) from tamoxifen treated lungs. (Ji) Low magnification view of the airways showing CCs and multiciliated cells. (Jii) shows a high magnification view of the distribution of $CC10^+$ cells around the boxed NEB. Note depletion of $CC10^+$ cells in the long term. Red arrowheads indicate remaining $CC10^+$ cells. (K) Thin sections (5μm) from lungs, 1 month post RBPJκ ablation stained for RBPJκ/FoxJ1/CGRP. Yellow arrowheads denote $RBPJκ^-FoxJ1^+$ cells. (L) Frequency of $RBPJκ^-FoxJ1^+$ cells associated with NEBs 1 month post RBPJκ ablation. Data represented as mean ± standard deviation, * denotes p <0.01, Student's t-test.

**Figure S6. Impact of RBPJκ depletion in multiciliated cells on CCs.** (A) Genetic strategy and tamoxifen treatment regimen in FoxJ1$^{creERTm/+}$; RBPJκ$^{flox/flox}$ to conditionally ablate RBPJκ in multiciliated cells. Thin sections from uninduced control (B) and tamoxifen treated lungs (C) stained with RBPJκ /CC10/FoxJ1 (Bi, Ci) and RBPJκ /FoxJ1 (Bii, Cii) . Note the loss of RBPJκ expression in multiciliated cells upon tamoxifen treatment. Yellow arrowheads denote the nucleus. (D-I) Phenotype of CCs post depletion of RBPJκ in multiciliated cells. Thin sections from uninduced control (D, F) and tamoxifen treated lungs (E, G) stained for CGRP/CC10/Scgb3a2 and CC10/Scgb3a2. Note that CCs are double positive for CC10 and Scgb3a2 around NEBs (D,E) and at terminal bronchioles (F,G). (H, I) Thin sections from uninduced control (H) and tamoxifen treated lungs (I) stained for CGRP/CC10/SPC. Note the lack of SPC expression in CCs upon RBPJκ ablation in multiciliated cells (white arrowheads).

**Figure S7. Impact of antibody mediated inhibition of Notch2/Notch1 and Jagged1/Jagged2 on *Uroplakin3a*$^{+}$ CCs.** (A-F) Lineage analysis of U-CCs (v-CCs). (A) Treatment regimen for labelling U-CCs and inhibition of Notch signalling using antibodies in Upk3a$^{CreERTm/+}$;Rosa$^{Tdtomato}$ mice. (B-D) Thin sections stained for CGRP/CC10/Tdtomato from Anti gp120 (B), Anti Notch2/Notch1 (C) and Anti Jagged1/Jagged2 (D) treated lungs. Yellow arrowheads indicate Tdtomato$^{+}$CC10$^{+}$ cells. Yellow dashed line demarcates NEB. (E, F) Thin sections stained for CGRP/Tdtomato/SPC from Anti Notch2/Notch1 (E) and Anti Jagged1/Jagged2 (F) treated lungs. Yellow arrowheads denote Tdtomato$^{+}$SPC$^{+}$ cells.

**SUPPLEMENTAL INFORMATION TITLES AND LEGENDS**

**Animal Handling**

Any procedure that could conceivably cause distress to the animals employed peri-procedure anaesthesia with isofluorane gas (Baxter Healthcare Corp.) delivered by an anaesthetic vaporizing machine in our animal facility. In addition, all animals were monitored for signs of distress and euthanized if in distress. Euthanasia was performed by anaesthetising the animals followed by cervical dislocation at inStem.

**Mouse strains**

All animal strains were housed under Specific Pathogen Free environment (SPF) conditions in the Animal Care and Resource Centre (ACRC) at the National Centre for Biological Sciences (NCBS). B6/J (C57BL/6), Scgb1a1$^{CreERTm/+}$ (C57BL/6), Rosa26-LSL-tdT (C57BL/6, Ai14), Upk3a$^{CreERT2}$ (C57BL/6), FoxJ1$^{CreERT2}$ (C57BL/6) and Rosa26-LSL-DTA (C57BL/6) strains were obtained commercially (The Jackson Laboratory). Floxed RBP-J strain was a kind gift from Dr. Mitsuru Morimoto, RIKEN. Frozen embryos of the Floxed RBP-J strain were obtained from RIKEN BRC, cryo-rederived at the Mouse Genome and Engineering Facility (MGEF) at NCBS and maintained on a C57BL/6 background. Scgb1a1$^{CreERTm/+}$ strain was crossed to the Floxed RBP-J strain to generate Scgb1a1$^{CreERTm/+}$; RBPJκ$^{flox/flox}$. FoxJ1$^{CreERT2}$ strain was crossed to Rosa26-LSL-DTA to generate FoxJ1$^{CreERT2}$; Rosa$^{DTA/+}$.

**Notch inhibition models**

<u>Pharmacological inhibition of Notch signalling by γ-secretase inhibitor - Dibenzazepine (DBZ)</u>. A stock solution of 92.6 mg/ml DBZ (Sigma) in Dimethyl Sulfoxide (DMSO) was prepared and stored at 4°C in the dark. For injections, the stock solution was diluted to 2.32 mg/ml working solution in MCT (0.5% w/v Methylcellulose, 0.1% Tween 80 in 0.9% NaCl solution). B6/J or Upk3a$^{creERT/+}$; Rosa$^{Tdtomato/+}$ (≥6 weeks of age) were injected intraperitoneally with DBZ (50μmol/kg) or MCT alone (vehicle control) every twelve hours for seven consecutive days and were sacrificed on the eighth day. Lungs were inflated with paraformaldehyde and fixed overnight 4°C and processed for embedding in paraffin. Thin sections were immunostained and imaged. Multi-channel images were used for cell analysis.

<u>Genetic ablation of RBPJκ in multiciliated cells</u>– FoxJ1$^{CreERT2}$;RBPJκ$^{flox/flox}$ mice (≥6 weeks of age) were injected with Tamoxifen (Sigma, 250mg/kg body weight) four times on alternate days before sacrificing at the timepoints indicated in the figures.

**Cell labelling for lineage analysis**

For labelling Upk3a-expressing cells, Upk3a$^{CreER/+}$; Rosa26$^{Tdtomato/+}$ heterozygotes ≥6 weeks of age) were injected with Tmx (Sigma, 250mg/kg body weight) three times on alternate days. For labelling all Club cells, Scgb1a1$^{CreERTm/+}$; Rosa26$^{Tdtomato/+}$ heterozygotes (≥6 weeks of age) were injected with Tmx (Sigma, 250mg/kg body weight) once. For lineage analysis post Notch inhibition, Upk3a$^{CreER/+}$; Rosa26$^{Tdtomato/+}$ or Scgb1a1$^{CreERTm/+}$; Rosa26$^{Tdtomato/+}$ animals were first induced with tamoxifen. After a 7-10 day tamoxifen wash out period, the induced animals were injected with Anti Notch1/Notch2 or Anti Jagged1/Jagged2 antibodies. They were then sacrificed at the timepoints shown schematically in the figures.

**Histology, Immunofluorescence, and Imaging**

Adult lungs were inflated with 1% (wt/vol) Paraformaldehyde + 2% low-melting agarose in PBS and immersed in the same fixative for 1hr at RT. The lobes were separated and left lobe was used for thick section analysis and the remaining lobes for immersed in the same fixative for another 4-5 hr at RT on a rocker. For thick sections (200 µm), the left lobe was embedded in 2% agarose and sectioned on a Compresstome. The sections were fixed again in 1% (wt/vol) Paraformaldehyde in PBS for 1hr at RT, washed with PBS and stored at 4°C. Rest of the other fixed lobes were incubated in PBS at 60°C overnight for melting the agarose and subsequently processed and embedded in paraffin for histological analysis. For Notch2ICD staining, lungs were inflated with 4% (wt/vol) Paraformaldehyde and incubated at 4°C overnight on a rocker. Fixed lungs were then washed with PBS five times, 5 min each at 4°C, processed and embedded in paraffin for histological analysis.

Heat mediated antigen retrieval was performed using antigen unmasking solution from Vector labs (pH9) prior to immunostaining. pH6 antigen unmasking solution was used when staining for Notch2ICD. Immunofluorescence analysis utilized the following antisera: Rabbit anti-Notch2ICD (abcam, 1:100 [thin sections]), Goat anti-Uteroglobin (Merk, 1:1000 [thin sections], 1:500 [thick sections]), Rabbit anti-CGRP (Sigma, 1:1000 [thin sections], 1:500 [thick sections]), Mouse anti-acetylated tubulin (Sigma, 1:1000 [thin sections], 1:500 [thick sections]), Goat anti-mUGRP1 (1:1000 [thin sections], 1:500 [thick sections]), Rabbit anti-Sftpc (Seven Hills Bioreagents, 1:500 [thin sections], 1:500 [thick sections]), Mouse anti-Foxj1 (eBioscience, 1:200), Mouse anti-RFP (Abcam, 1:300), Rabbit anti-RFP (Rockland, 1:300), Hamster anti-Pdpn (Developmental Studies Hybridoma Bank, 1:250), Rabbit anti-Nkx2-1 (Abcam, 1:500), Mouse anti-Foxi1 (Origene, 1:100), Rabbit anti-DCLK1 (Abcam, 1:200), Rabbit anti-Δp63 (Cell Signalling Technologies, 1:200), Mouse anti-Muc5Ac (Thermofisher Scientific, 1:200), Rabbit anti-RBPUSH (Cell Signalling Technologies, 1:200), Rabbit anti-ABCA3 (Seven Hills Bioreagents, 1:500), Mouse anti-Sox2 (Santa Cruz, 1:50), Rabbit anti-

TTF1 (Abcam, 1:100), Goat anti-CGRP (Abcam, 1:500), Rabbit anti-Acetylated tubulin (Abcam, 1:1000). Alexa 405/488/568/647-conjugated Donkey anti-mouse/rabbit/goat (Invitrogen, 1:300). For thick sections, 0.5% Triton-X-100 in 1X PBS was used for preparing dilutions for all antibodies. Sections were imaged either on Zeiss LSM-780 or LSM-980 or Olympus FV3000 4/5 lasers, laser-scanning confocal microscopes. For thick sections, z-stacks were collected at 3-5µm intervals and maximum intensity projection in the z-dimension were created for 3-D reconstruction of the airways.

**Quantification of cell frequencies**

Cell frequencies were calculated from immunostained paraffin sections (4-5µm). Multi-channel images were acquired using confocal microscopes and cell frequencies were obtained by manual counting in images using Fiji (ImageJ). Nuclei were identified by the presence of DAPI and cells that were positive for the respective markers were counted. The total length of the airway was determined by drawing a line that traces the length of the airway. The frequencies were then calculated as number of cells per mm of airway and the frequencies were plotted in Microsoft Excel. In our analysis, NEB associated cells were identified as nuclei that were in direct contact with $CGRP^+$ cells. Closed airways with a diameter of ≤200 µm or 200 µm from the BADJ of open-ended airways that transition into the alveoli were considered as terminal bronchioles. In thick sections, NEBs were identified as clusters of >2 $CGRP^+$ cells. All frequency plots are either totals in pie charts or averages ± standard deviation in bar plots. At least 2 thick and 2 thin sections from 3 animals were analysed per condition. Student’s t-test was used for calculating the significance in the statistical analysis.

### Figure S1

**Impact of antibody mediated inhibition of Notch2/Notch1 and Jagged1/Jagged2 on CCs.**

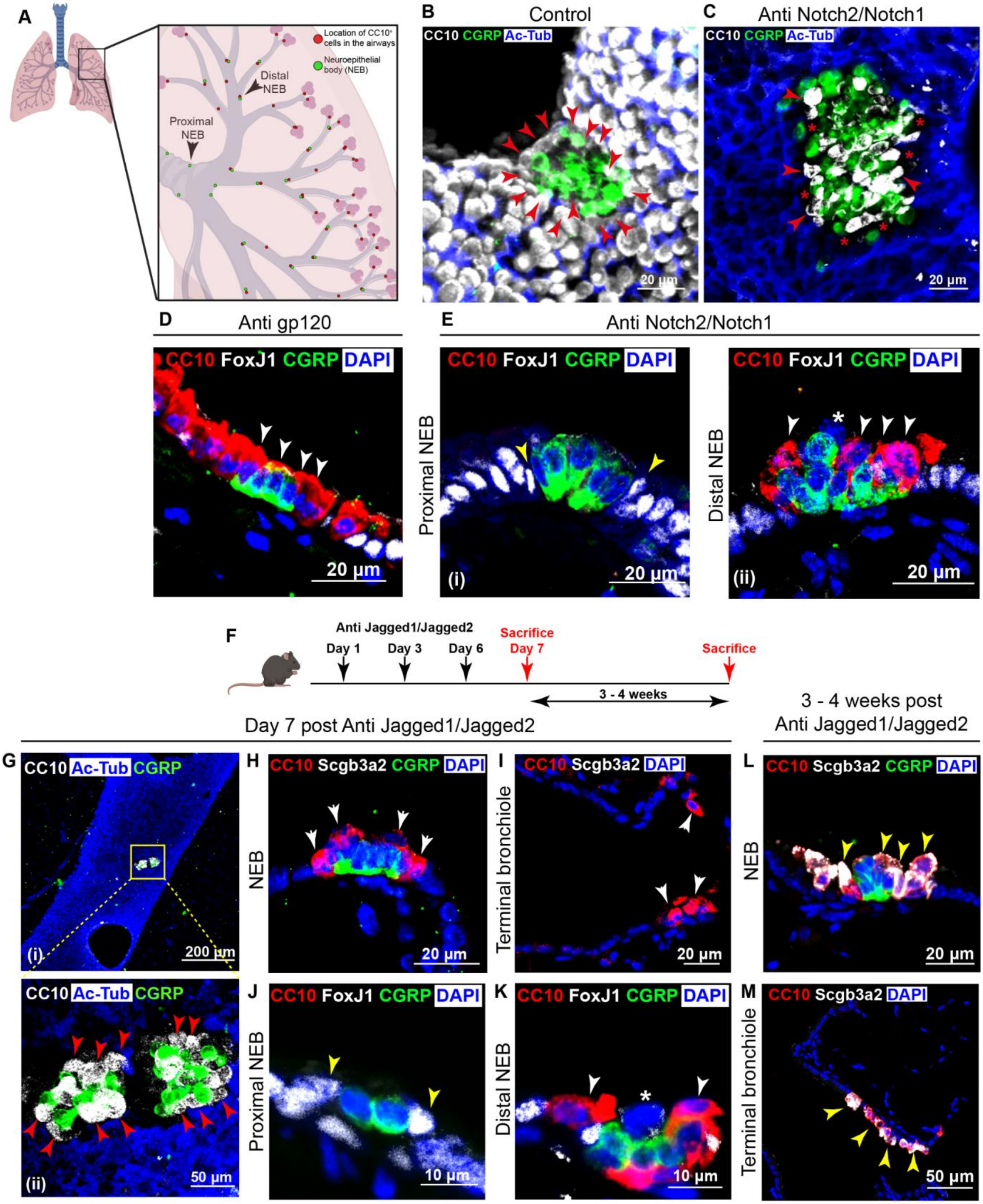

Figure S2

**CCs surrounding NEBs retain airway identity but do not differentiate into other known airway or alveolar cell states upon antibody mediated inhibition of Notch2 and Notch1.**

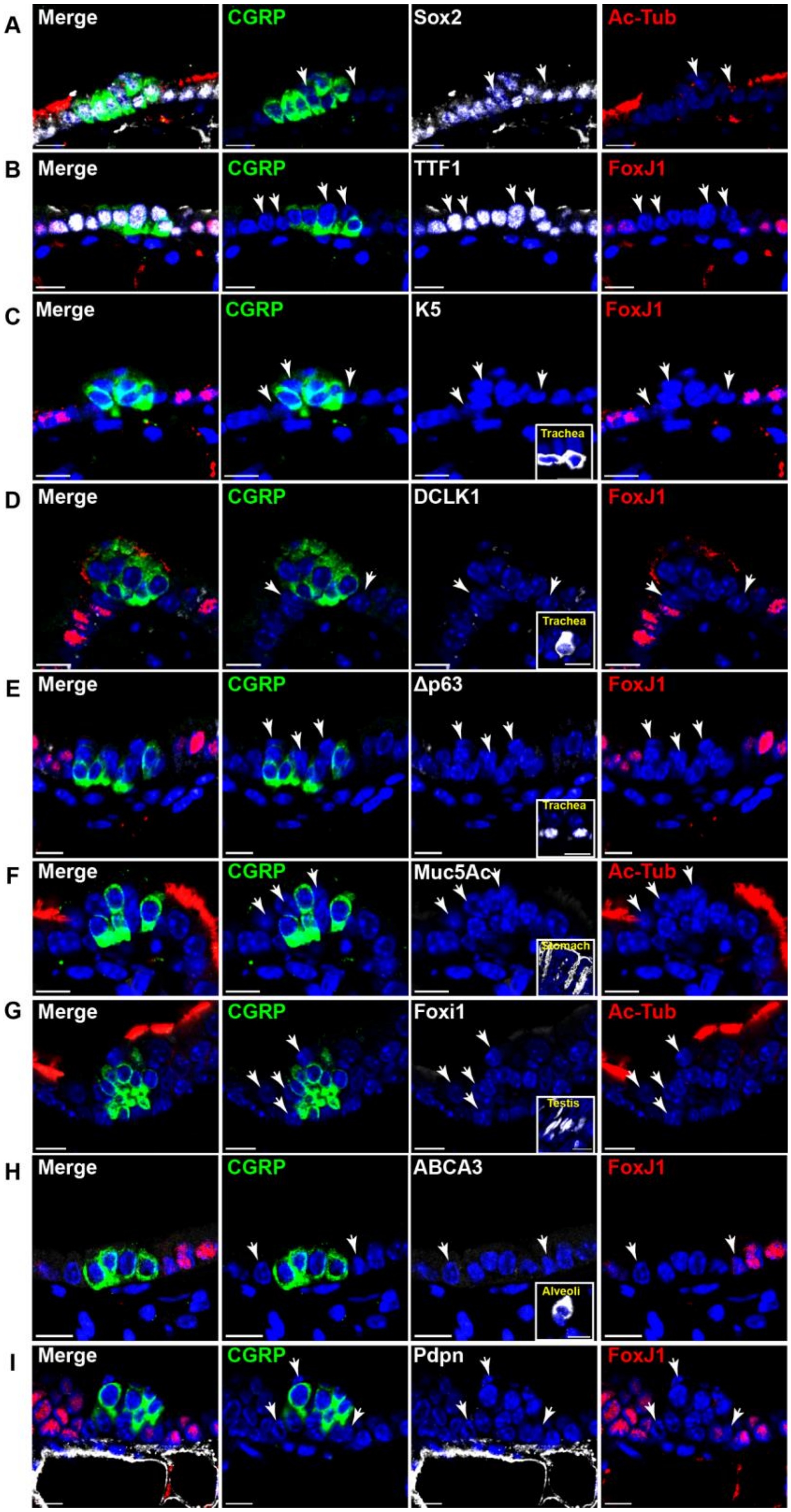

Figure S3

**CCs surrounding NEBs and at terminal bronchioles express the alveolar type II marker - Surfactant protein C (SPC) upon antibody mediated inhibition of Jagged1 and Jagged2.**

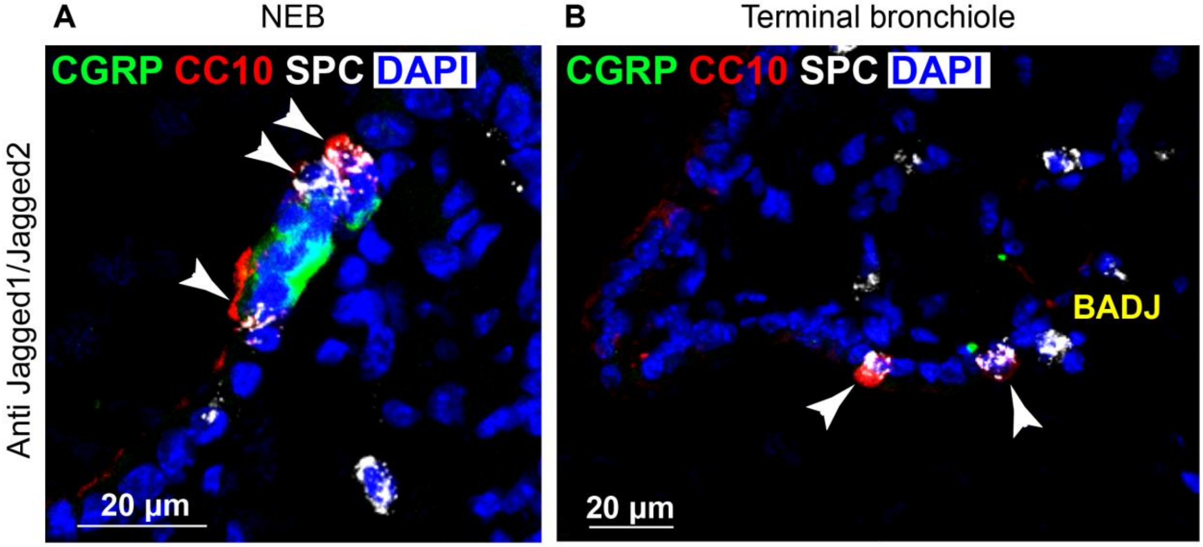

Figure S4

**Impact of small-molecule mediated (γ-secretase inhibitor - Dibenzazepine (DBZ)) inhibition of Notch signaling on CCs.**

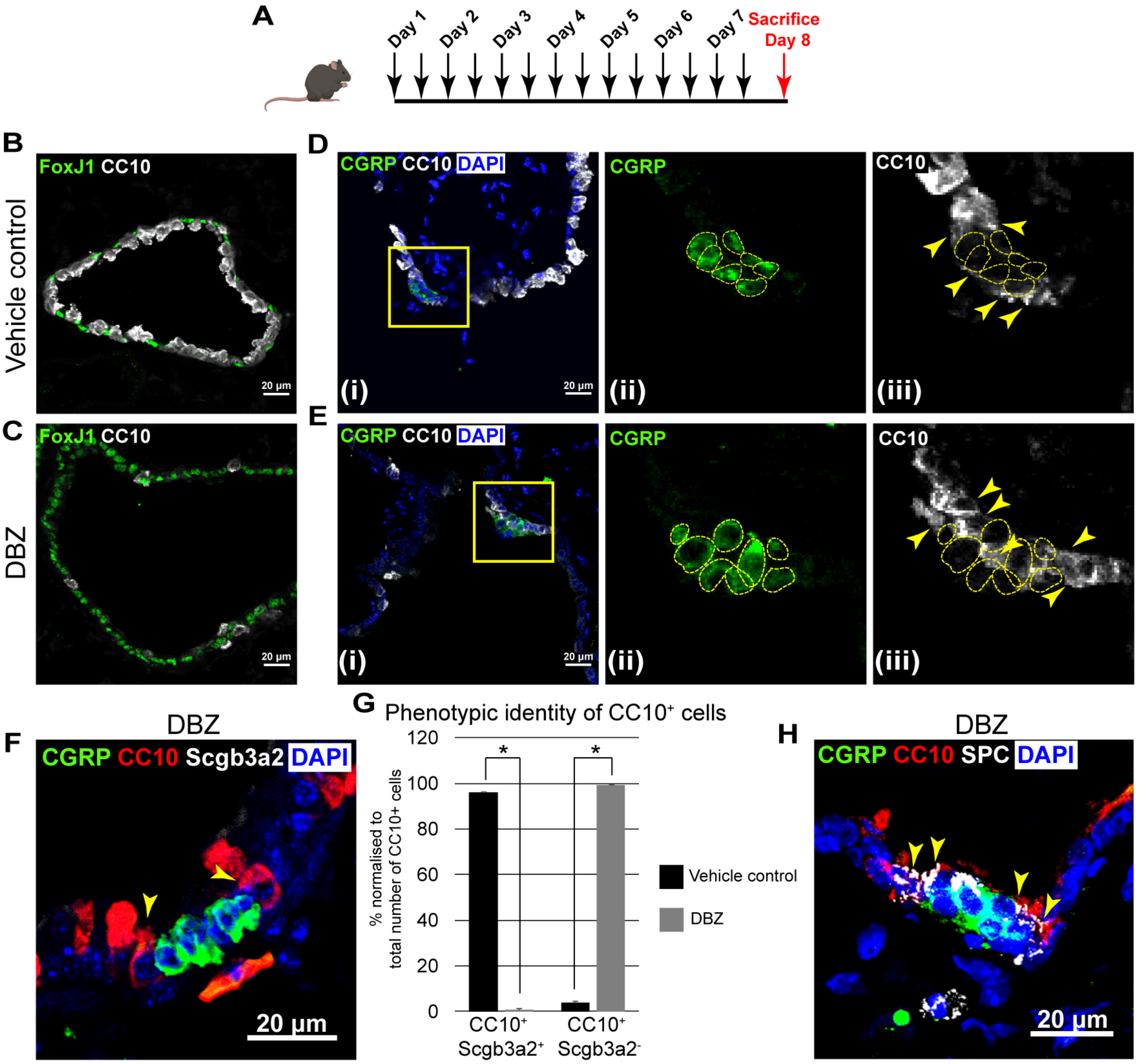

Figure S5

**Impact of RBPJκ depletion on CCs.**

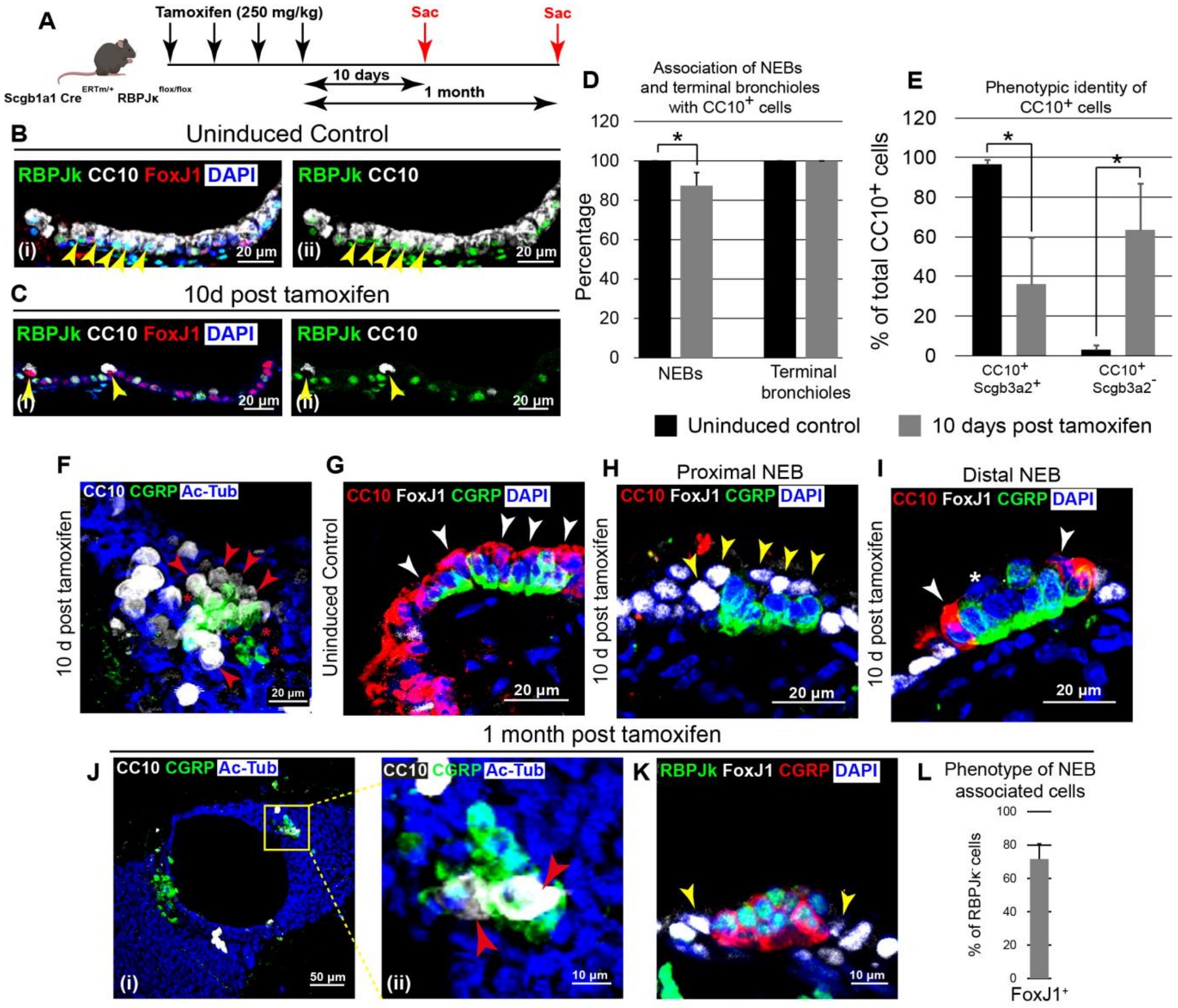

### Figure S6

**Impact of RBPJκ depletion in multiciliated cells on CCs.**

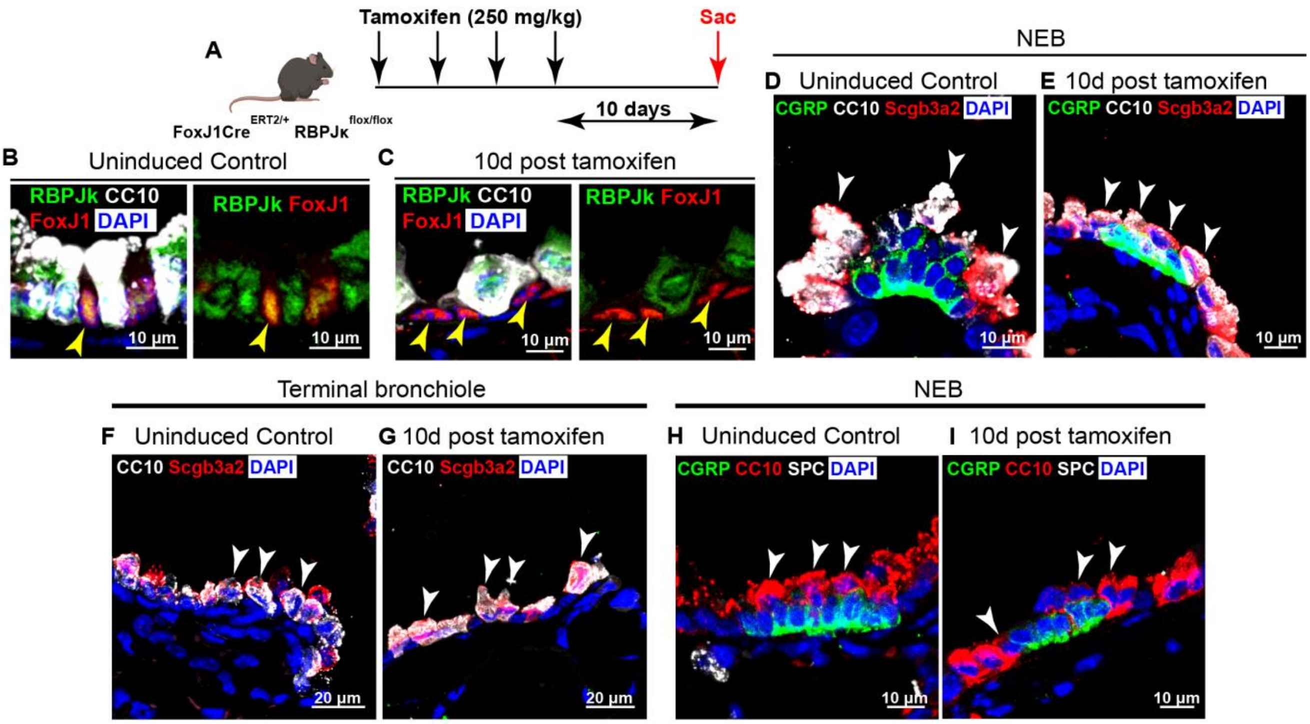

Figure S7

#### Impact of antibody mediated inhibition of Notch2/Notch1 and Jagged1/Jagged2 on *Uroplakin3a*⁺ CCs

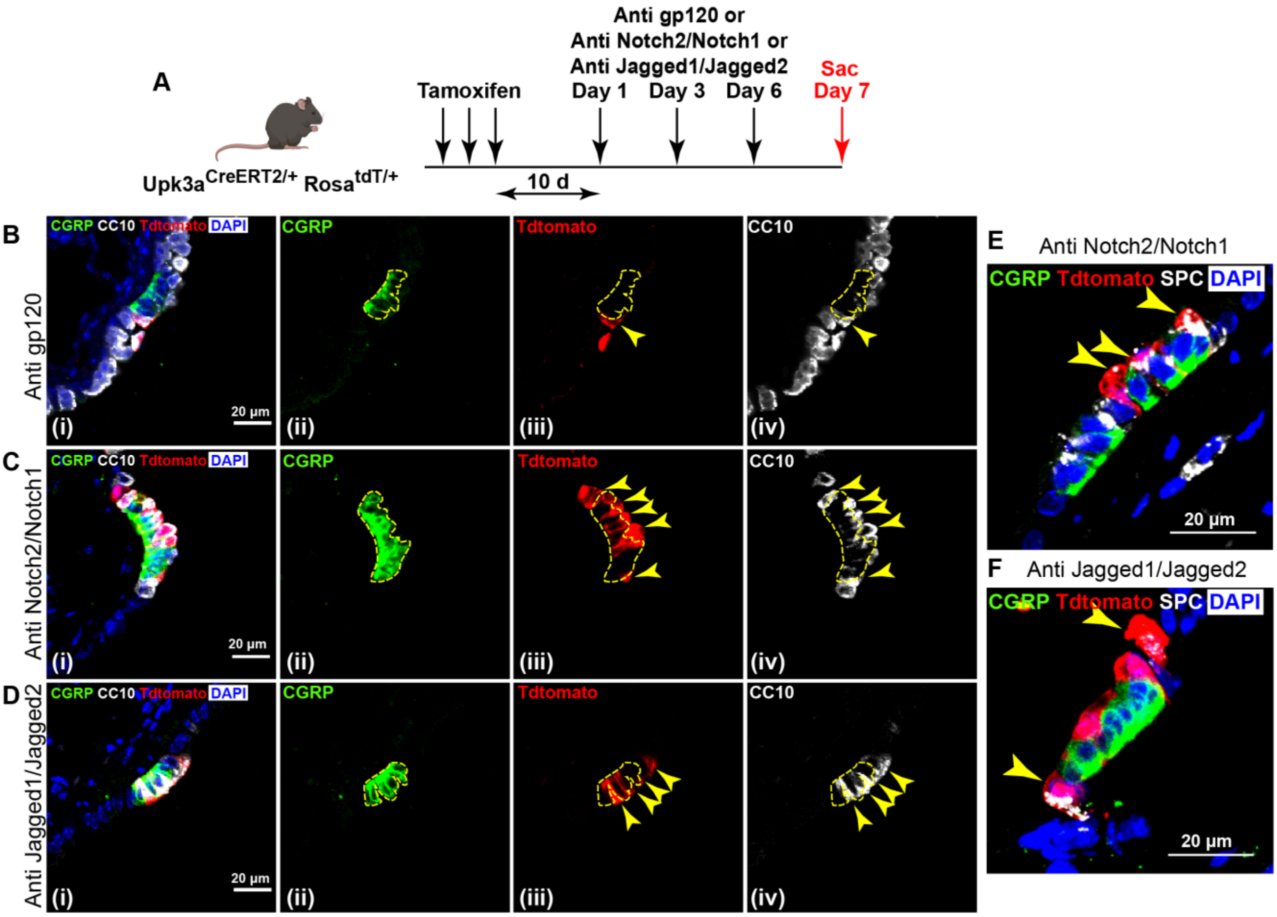
